# Organism-Wide Analysis of Sepsis Reveals Mechanisms of Systemic Inflammation

**DOI:** 10.1101/2023.01.30.526342

**Authors:** Michihiro Takahama, Ashwini Patil, Katherine Johnson, Denis Cipurko, Yoshimi Miki, Yoshitaka Taketomi, Peter Carbonetto, Madison Plaster, Gabriella Richey, Surya Pandey, Katerina Cheronis, Tatsuki Ueda, Adam Gruenbaum, Steven M. Dudek, Matthew Stephens, Makoto Murakami, Nicolas Chevrier

## Abstract

Sepsis is a systemic response to infection with life-threatening consequences. Our understanding of the impact of sepsis across organs of the body is rudimentary. Here, using mouse models of sepsis, we generate a dynamic, organism-wide map of the pathogenesis of the disease, revealing the spatiotemporal patterns of the effects of sepsis across tissues. These data revealed two interorgan mechanisms key in sepsis. First, we discover a simplifying principle in the systemic behavior of the cytokine network during sepsis, whereby a hierarchical cytokine circuit arising from the pairwise effects of TNF plus IL-18, IFN-γ, or IL-1β explains half of all the cellular effects of sepsis on 195 cell types across 9 organs. Second, we find that the secreted phospholipase PLA2G5 mediates hemolysis in blood, contributing to organ failure during sepsis. These results provide fundamental insights to help build a unifying mechanistic framework for the pathophysiological effects of sepsis on the body.

## INTRODUCTION

Molecules, cells, and tissues with immune functions are ubiquitous in the body. While the organismal nature of the immune system is vital for the host against infection, the systemic dysregulation of immune processes in response to infectious and non-infectious triggers can be harmful. For example, sepsis is a systemic host response to infection with life-threatening consequences.^1, 2^ The disease is a global health issue in need of targeted therapies addressing the short- and long-term effects on the host.^3–7^ Our knowledge of the mechanisms underlying the impact of sepsis on the body is rudimentary, as highlighted by expert consensus in the field of sepsis.^8^ The timing and location of events that take place across organs other than blood during sepsis remain unclear. Sepsis is thus a clear example for which learning the multifactorial effects of the disease on the molecules, cells, and tissues of the whole body is critically important for basic and clinical sciences.

A myriad of cells and molecules has been linked to sepsis. Numerous studies have established immune and endothelial cells together with cytokines and the complement and coagulation systems as key cellular and molecular factors in the pathogenesis of sepsis.^9–12^ However, the links between the molecular and cellular factors that produce the damaging impact of sepsis for the body have not been systematically mapped. For example, the uncontrolled, systemic activity of cytokines contributes to tissue injury and organ failure,^13^ but it is unclear which cytokines – alone or in combination – impact which cells and tissues across the body. As a result, we lack a unifying framework to understand how the cytokine network functions in sepsis, including the network’s hierarchy, interactions, and feedback loops.^14^

In addition, many types of cells die or divide at abnormal rates during sepsis.^9, 15, 16^ The number of lymphocytes drops,^17, 18^ while that of neutrophils surges during sepsis,^19^ contributing to the negative effects of the disease on the immune system of survivors.^9, 16, 20^ However, we have a limited understanding of which molecules are responsible for the effects of sepsis on immune and non-immune cells across various tissue contexts.^8^ Therefore, to better understand the systemic effects of sepsis, we must build a mechanistic framework explaining the causal relationships between the key molecular and cellular factors of the disease at the level of the whole organism.

Here we used mouse models of sepsis to obtain a dynamic, organism-wide map of the pathogenesis of the disease, revealing the spatial and temporal patterns of both known and previously unrecognized effects of sepsis on the body. Our work has identified a cytokine circuit arising from the pairwise effects of TNF plus IL-18, IFN-γ, or IL-1β, which recapitulates a large fraction of the effects of sepsis across organs. Going further, by mapping the effects of these three cytokine pairs on the abundance of 195 cell types across 9 organ types, we provided a mechanistic basis for the effects of the disease on cells across tissues. Lastly, by mining our organism-wide maps of sepsis for candidate regulators of the disease, we discovered an interorgan pathway detrimental to the host whereby the release from the gut into the blood circulation of a secreted phospholipase, PLA2G5, led to hemolysis and the failure of multiple organs.

## RESULTS

### Organism-Wide Analysis of Gene Expression Changes in Mouse Models of Sepsis

To study the dynamics of gene expression changes induced by systemic inflammation across the body, we used two mouse models of sepsis (**Figure 1A**). We used the intra-peritoneal injection of lipopolysaccharide (LPS) at sublethal (5 mg/kg) and lethal (15 mg/kg) doses to mimic systemic bacterial inflammation (**Figure 1B**).^21^ In addition, we used cecal ligation and puncture (CLP), which is considered the gold standard model for sepsis due to its high clinical relevance.^22, 23^ We used CLP to trigger a polymicrobial infection starting in the abdominal cavity and ranging in disease severity based on the position of the cecal ligation and the number and size of cecal punctures (**Methods**) (**Figure 1B**). First, we performed whole-tissue RNA-seq across a dozen organ types upon sublethal LPS injection (**Figure 1A**).^24, 25^ We profiled 13 tissues per animal, including bone marrow, brain, colon, heart, inguinal lymph node (iLN), kidney, liver, lung, peripheral blood mononuclear cells (PBMCs), skin, small intestine, spleen, and thymus, at six time points after LPS injection (0.25, 0.5, 1, 2, 3, and 5 days) – to observe early and late phases of the disease – and from untreated control mice using four biological replicates per condition. In total, we identified 10,003 genes that were differentially expressed between LPS-injected and control samples across one or more tissues and displayed both intra- and cross-tissue expression patterns (**Figures 1C** and **Table S1A**).

**Figure 1.**
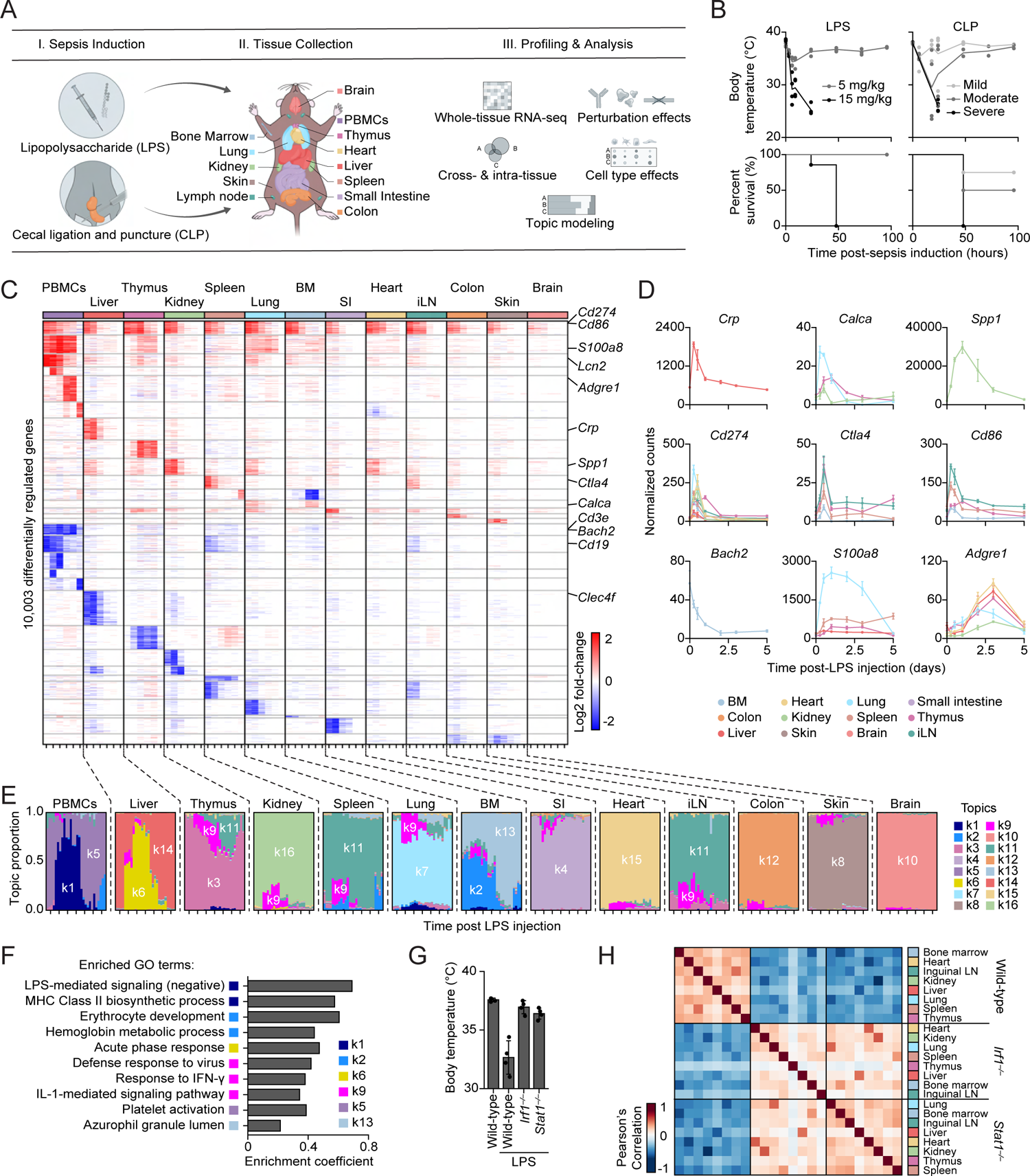
Whole-Tissue Gene Expression Reveals the Molecular Effects of Sepsis across Organs. (A) Schematic overview of the study design. PBMCs, peripheral blood mononuclear cells. (B) Measurements of rectal temperature (top) and survival (bottom) after LPS injection at sublethal (5 mg/kg) or lethal (15 mg/kg) doses (left, n = 4), or cecal ligation and puncture (CLP) surgeries leading to mild, moderate, and severe sepsis (Methods) (right, n = 5). (C) Heatmap of differentially expressed genes (rows) from whole-tissue mRNA profiles ordered by k-means clustering (horizontal lines), organ types (top, colors) and times (bottom, tick marks for 0.25, 0.5, 1, 2, 3, and 5 days) post-sublethal LPS injection. Values are log2 fold-changes relative to matching, untreated organ (FDR-adjusted p-value < 0.01; absolute fold change > 2; n = 2-4 for PBMCs and 3-4 for other organs). (D) Normalized counts for indicated genes and organs (color). BM, bone marrow; iLN, inguinal lymph node; Error bar, SEM (n = 3-4). (E) Structure plot of the estimated membership proportions for a topic model with k = 16 topics (colors) fit to 364 tissue samples across 13 organ types (top) from LPS-injected mice (Methods). Each vertical bar shows the cluster membership proportions for a single tissue sample ordered over time (bottom, tick marks for 0, 0.25, 0.5, 1, 2, 3, and 5 days post-sublethal LPS injection) for each organ type. SI, small intestine. (F) Pathway enrichment analysis using differentially expressed genes in each topic from E. Shown are enrichment coefficients (x axis) for indicated Gene Ontology (GO) terms (y axis). (G) Measurements of rectal temperature in mice of indicated genotypes at 12 hours post-sublethal LPS injection (5 mg/kg) or in untreated controls. Error bars, SD (n = 4). (H) Similarity (Pearson’s correlation coefficient) between whole-tissue mRNA profiles from mice of indicated genotypes at 12 hours post-sublethal LPS injection (5 mg/kg). **See also** Figure S1 and Table S1.

Our data revealed the spatiotemporal expression patterns of numerous molecules and processes associated with sepsis and systemic inflammation. For example, several mRNAs encoding common clinical biomarkers were upregulated over time such as *Crp* (liver), *Calca* (lung and kidney, early, and thymus, late), and *Spp1* (kidney) (**Figure 1D**). The expression of *Cd274* (PD-L1), *Ctla4*, and *Cd86*, encoding membrane-bound, costimulatory proteins thought to play a role in the immune deficiencies observed in sepsis,^9, 26^ was upregulated across numerous tissues: 12/13 for *Cd274*, 6/13 for *Cd86*, and 3/13 for *Ctla4* (**Figure 1D**). Changes in immune cell population dynamics were reflected by the expression patterns of immune cell marker genes (**Figure 1D** and **S1A**), such as *Bach2* for erythropenia,^27, 28^ *S100a8* neutrophil accumulation in lungs,^19^ *Cd3e* and *Cd19* for T and B cell lymphopenia,^29^ and *Adgre1* for multi-tissue accumulation of macrophages.^30^ Going further, to systematically examine how sepsis biomarkers varied in expression, we examined the expression patterns of 258 genes encoding biomarkers associated with sepsis in the literature.^31^ We observed changes in sepsis biomarker expression across all organs, with a range of effects including the lowest in brain with 9.7% (25/258) of biomarker genes regulated to the three highest in lung, thymus, and PBMCs with 37.5% (97/258), 36.7% (95/258), and 35.5% (92/258) of biomarker genes regulated, respectively (**Figure S1B**). Interestingly, when examining total changes in gene expression, we found that non-lymphoid tissues returned to their transcriptional steady state within five days of LPS injection, whereas lymphoid tissues did not (**Figure S1C**), which is reminiscent of the reported link between sepsis and long-term immune defects.^20, 32^ Lastly, we compared the organism-wide effects of LPS to those obtained using the CLP model of sepsis. We found a high degree of similarity between the whole-tissue gene expression profiles of LPS and severe CLP (**Figure S1D** and **Table S1C**), including overlaps in the identity of regulated genes ranging from 29.5% in heart to 68% in thymus (**Figure S1E**). In addition, the changes in expression in sepsis biomarkers and immune cell markers were similar in LPS and CLP (**Figure S1F**), suggesting that using the LPS model for further mechanistic studies is likely to yield ©nsights that also apply to the CLP model.

Next, we analyzed the LPS time series data using grade of membership models to examine the impact of sepsis on intra- and cross-tissue states. Grade of membership models, also known as topic models, cluster samples by allowing each sample to have partial membership in multiple biologically distinct clusters or “topics”,^33^ as opposed to traditional clustering methods that assign a sample or a gene to a single cluster. We first fit the grade of membership model to our LPS data using 16 topics, and generated structure plots of estimated membership proportions for all 364 whole-tissue RNA-seq profiles encompassing 13 tissues and 6 time points post-LPS injection in addition to control, untreated samples (**Figure 1E**). Second, to determine which genes and processes explain each topic, we used the quantitative estimates of the mean expression of each gene in each topic as provided by the grade of membership models to perform gene set enrichment analyses (**Methods**). Several topics reflected the known biology of all the tissues profiled, such as topics k4, k7, and k15, which reflected basic functions of the small intestine, lungs, and heart, respectively (**Table S1D**). Other topics captured processes driven by LPS-induced sepsis as highlighted by dynamic changes in the levels of membership of tissue samples to those topics over time (**Figure 1E-F**). For example, some topics reflected responses linked to an influx of granulocytes in PBMCs, which is linked to clinical deterioration,^34^ and, to a lesser extent, in lungs and bone marrow (k1), or to acute inflammatory response of the liver (k6). Other topics captured changes in cell populations such as erythropenia in the bone marrow (k2) and neutrophil proliferation and recruitment in the spleen and lungs (k13). Lastly, topic k9 reflected organism-wide changes in interferon-stimulated genes (ISGs). To investigate the transcriptional regulation of ISGs across tissues, we compared the effect of LPS on whole-tissue gene expression between wild-type, *Irf1^−/−^*, and *Stat1^−/−^* mice. We focused on the transcription factors (TFs) IRF1 and STAT1 because: (i) they were predicted computationally to be the TFs regulating the genes driving cluster K9 (**Figure S1G**); (ii) they are among the few TFs in topic k9 which are upregulated across all 13 organs profiled upon LPS injection (**Figure S1H**); and (iii) even though they are well-known regulators of ISGs,^35^ the target genes of these two TFs across tissues have not been mapped. We found that a sublethal injection of LPS in *Irf1^−/−^* and *Stat1^−/−^* mice did not lead to the decrease in body temperature seen in wild-type mice (**Figure 1G**), in agreement with published data,^36, 37^ and abrogated the multi-tissue ISG response driving topic k9 (**Figure 1H** and **S1I-J**). The LPS-induced genes regulated by IRF1 and STAT1 across organs overlapped for the most part, although some tissue-specific effects were also observed, such as in liver, kidney, and lung (**Figure S1J**).

### The Pairwise Effects of TNF plus IL-18, IFN-γ, or IL-1β Collectively Recapitulate Most of the Transcriptional and Physiological Effects of Sepsis

To link the intra- and cross-tissue effects of sepsis to specific molecules, we first investigated the effects of the following six cytokines: IFN-γ, IL-1β, IL-6, IL-10, IL-18, and TNF, which are key systemic factors involved in sepsis and cytokine storm syndromes.^9, 13^ All six cytokine mRNAs and proteins were respectively upregulated across tissues and in the blood within hours of LPS injection (**Figure 2A**). We hypothesized that comparing the organism-wide changes in gene expression induced by LPS versus recombinant cytokines injected intravenously would help to reveal (1) how much of the effects of LPS sepsis can be explained by interorgan cytokine signaling, and (2) which cytokines impact the states of which tissues, pathways, and cell types. To test this idea, we systematically measured the effects of the six selected cytokines alone or in pairwise combinations (15 pairs) by performing whole-tissue RNA-seq at 16 hours post-cytokine injection on the 9 tissues capturing most of the changes induced by sepsis across the body (*i.e.*, bone marrow, colon, heart, inguinal lymph node, kidney, liver, lung, spleen, and thymus). All cytokine singles and pairs led to significant changes on tissue states, ranging from 14 (IL-10) to 431 (IL-1β) for singles and 12 (IL-6 + IL-10) to 7,083 (TNF + IL-18) for pairs in numbers of total differentially expressed genes between treated and control tissues across all 21 cytokine singles and pairs tested (**Figure S2A-B**). Next, we measured the overlap between differentially expressed genes driven by LPS and single and pairwise recombinant cytokines. Out of all 21 conditions tested (6 singles and 15 pairs), we found a striking agreement between the genes regulated by LPS and three cytokine pairs: TNF with IL-18 (14.9 in LN to 56.8% in kidney), IFN-γ (3.6 in LN to 38.2% in thymus), or IL-1β (1.9 in LN to 28.2% in thymus) (**Figure 2B-C**). Consistent with these results, the transcriptional effects of injecting©e mice with plasma from LPS-injected mice overlapped well with LPS effects (4.74% in colon to 20.5% in liver) (**Figure 2B**). These similarities in tissue transcriptional states between LPS and the three cytokine pairs TNF plus IL-18, IFN-γ, or IL-1β also reflected a significantly higher proportion of the 258 sepsis biomarker genes being regulated by these pairs, 45.7% (118/258), 43.8% (113/258), 32.5% (84/258), respectively, compared to the other 12 pairs tested (8.1% ± 7.3 SD) (**Figure S2C**). Furthermore, the overlap in tissue expression profiles between these three cytokine pairs was also comparable to those of CLP and viral sepsis induced by Vaccinia virus strain Western Reserve (WR) (**Figure 2D**).^24^

**Figure 2.**
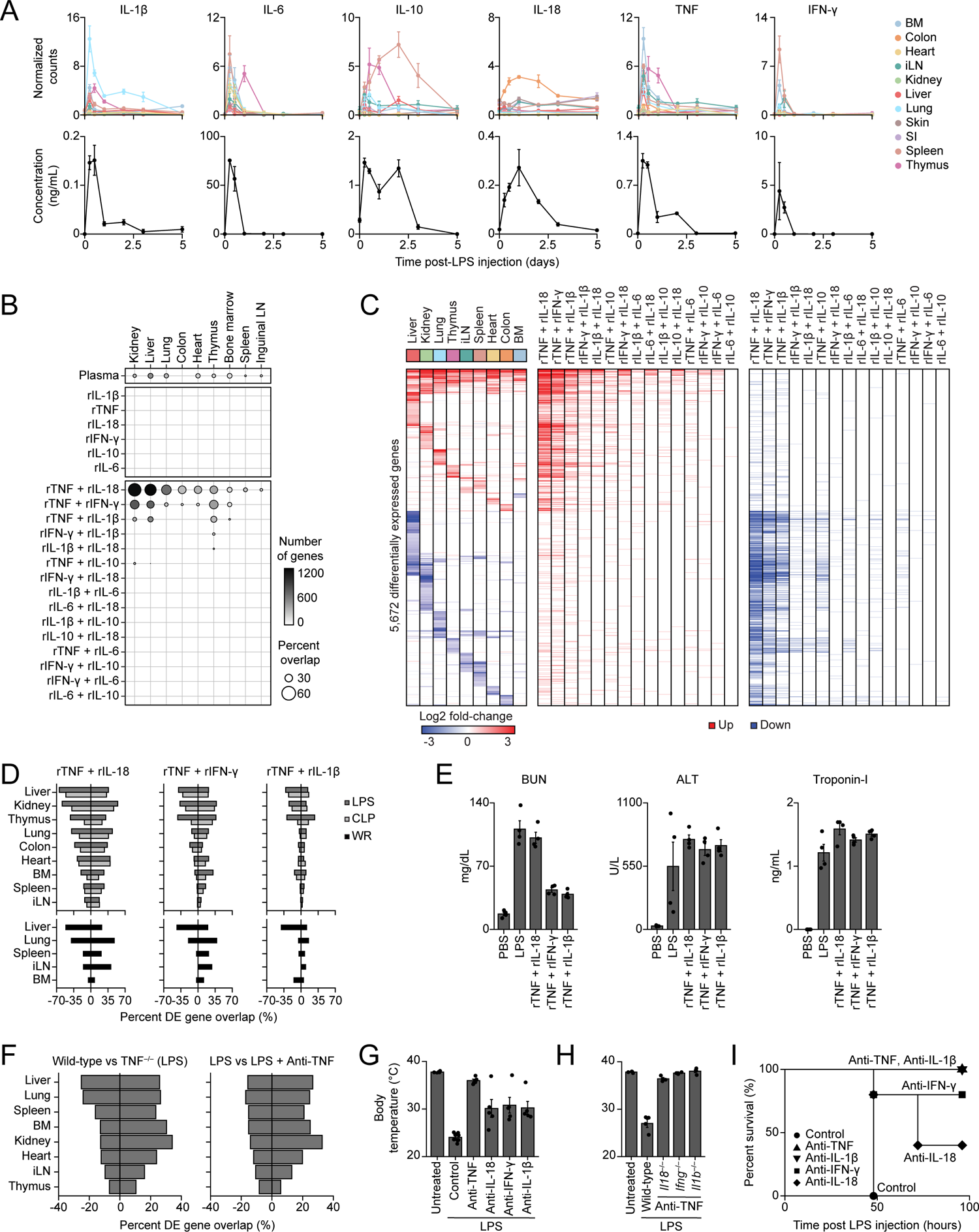
The Pairwise Effects of TNF plus IL-18, IFN-γ, or IL-1β Recapitulate Most of the Transcriptional and Physiological Effects of Sepsis. (A) Normalized counts (top) and blood concentration (bottom) for indicated cytokine genes and proteins upon sublethal LPS injection. Color, organ; BM, bone marrow; iLN, inguinal lymph node; SI, small intestine; Error bars, SEM (n = 3-4). (B) Percentages (circle) and numbers (color scale) of genes differentially expressed upon injection with indicated recombinant cytokines (rows) across organs (columns) that match the genes regulated by sublethal LPS at 12 hours post-sepsis induction. Plasma, naive mice injected with plasma fr©LPS-injected mice. (C) Heatmap (left) of differentially expressed genes (rows) from whole-tissue mRNA profiles ordered by k-means clustering and organ types (top, colors) at 12 hours after sublethal LPS injection. Values are log2 fold-changes relative to matching, untreated organs (FDR-adjusted p-value < 0.01; absolute fold change > 2; n = 4). Genes up- or down-regulated by indicated recombinant cytokine pairs in at least of one of the 9 tissues profiled are indicated in red and blue, respectively. (D) Percentages (x axis) of genes differentially expressed in tissues (rows) upon injection of the indicated three cytokine pairs that match the genes regulated by bacterial (LPS, CLP; top) or viral (WR; bottom) sepsis. WR, Vaccinia virus©rain Western Reserve. (E) Serum levels of indicated organ injury markers at 24 hours post-injection of a sublethal LPS dose or PBS as control, or 12 hours post-injection of indicated recombinant cytokine pairs. Error bars, SEM (n = 4). (F) Percentages (x axis) of genes differentially expressed in tissues (rows) upon sublethal LPS injection in mice treated with anti-TNF antibodies (1 h prior to LPS; left) or *Tnf^−/−^* mice (right) that match the genes regulated by LPS in wild-type, untreated mice. (G-H) Measurements of rectal temperature in mice of indicated genotypes with or without indicated neutralizing antibody pre-treatment at 24 hours post-lethal (G) or 12 hours post-sublethal (H) LPS injection.or bars, SD (n = 5-10). (I) Survival curves of mice injected with a lethal dose of LPS with or without indicated neutralizing antibody pre-treatment (n = 5). **See also** Figure S2 and Table S2.

To assess the physiological effects of the three cytokines pairs of interest on the body, we measured the blood levels of tissue injury markers and found that, as in LPS, all three pairs led to a significant increase in kidney, liver, and heart injury markers (**Figure 2E**). Furthermore, TNF plus IL-1β displayed a dose-dependent relationship between the quantity of cytokines administered and the body temperature and survival of the host (**Figure S2D**). Next, we investigated the impact of perturbing these four cytokines individually or pairwise using genetic deletions and neutralizing antibodies during sublethal and lethal LPS sepsis. First, we found that TNF deletion or neutralization strongly decreased the number of genes regulated by LPS sepsis across tissues, ranging from 9.2 (knockout) and 6.6% (blockade) in thymus to 25.5% (knockout) in liver and 23% (blockade) in kidney (**Figure 2F**). Second, neutralizing antibodies against IL-18, IFN-γ, or IL-1β all rescued mice injected with a lethal dose of LPS from a severe body temperature drop, albeit to a lesser extent than blocking TNF alone which sufficed to completely prevent temperature loss (**Figure 2G**). *Il18^−/−^*, *Ifng^−/−^* or *Il1b^−/−^* mice injected with anti-TNF neutralizing antibodies kept body temperatures at steady state levels upon LPS challenge (**Figure 2H**). Third, blocking TNF or IL-1β led to 100% survival in mice challenged with a lethal dose of LPS, whereas blocking IL-18 or IFN-γ led to partial survival (**Figure 2I**). Therefore, the similarities in tissue transcriptional states between sepsis and the three cytokine pairs reflect similarities in physiological effects, including tissue injury, body temperature, and survival.

Lastly, we compared the tissue transcriptional states induced by cytokine pairs to their composite single cytokines to classify the pairwise effects of each pair across organs as synergistic, antagonistic, or additive. Out of the 15 cytokine pairs tested, we found that TNF plus IL-18, IFN-γ, or IL-1β regulated the most genes across all organs with 7,083, 4,071, and 2,452 total regulated genes, respectively, compared to an average gene number of 382 ± 298 SD for the other 12 pairs tested (**Figure 3A**). The high numbers of regulated genes by the top three cytokine pairs compared to their matching singles were driven by strong synergistic and antagonistic regulatory events across all organs tested (**Figure 3B-D** and **S3A-B**). For example, liver displayed some of the highest proportions of synergistic and antagonistic genes across all three cytokine pairs: 10.2 and 30.3% for TNF + IL-18, 6.8 and 15.9% for TNF + IFN-γ, and 2.7 and 8.7% for TNF + IL-1β, respectively, whereas bone marrow displayed the lowest numbers of synergistic and antagonistic genes (2.1 to 11.2% across all three pairs) (**Figure 3C**). Interestingly, these three cytokine pairs regulated genes showing both shared and pair-specific patterns of expression, as shown in the examples of liver and kidney (mostly shared), and heart, spleen, and LN (mostly pair-specific) (**Figure 3D and S3B**). A gene-centric analysis revealed that, similarly to LPS and CLP, cytokine pairwise effects impacted multiple sepsis biomarker genes (**Figure 3E-H**). Taken together, these results supported a model whereby three cytokine pairs interact in each tissue to yield transcriptional and physiological states that closely resemble those induced by sepsis.

**Figure 3.**
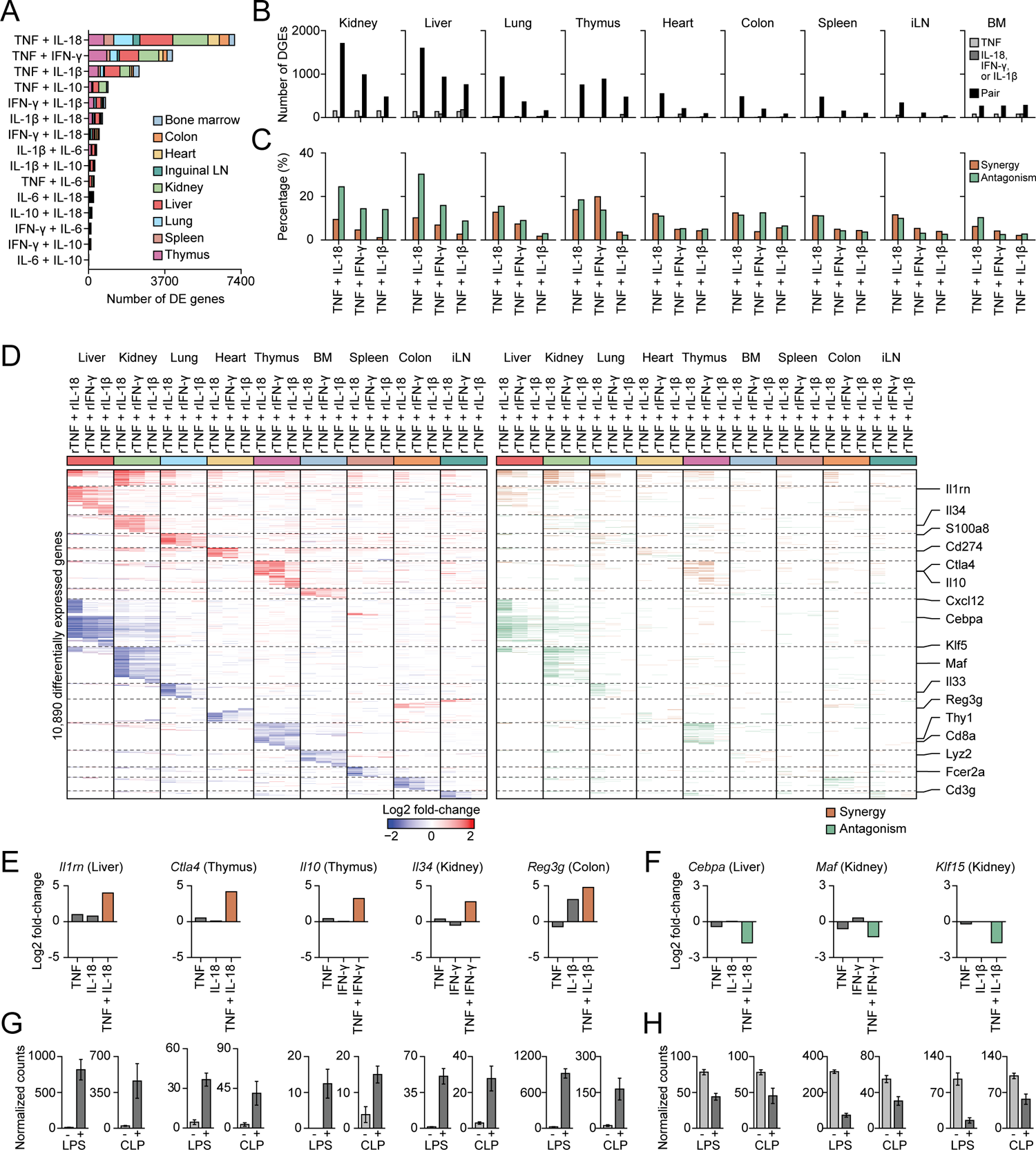
The Cytokine Pairs Composed of TNF plus IL-18, IFN-γ, or IL-1β Yield Nonlinear Effects on Tissue Transcriptional States. (A) Numbers of genes (x axis) regulated by indicated cytokine pairs (rows) but not by matching single cytokines. Color, organ type; LN, lymph node. (B) Numbers of genes (y axis) regulated in each tissue type by indicated cytokine p©s and composite singles. (C) Percentages (y axis) of differentially expressed genes in each tissue type displaying synergistic (orange) or antagonistic (green) in indicated cytokine pairs relative to matching single cytokines. (D) Heatmap (left) of differentially expressed genes (rows) from whole-tissue mRNA profiles ordered by k-means clustering and organ types (top, colors) at 16 hours after injection of indicated recombinant cytokine pairs. Values are log2 fold-changes relative to matching, untreated organs (FDR-adjusted p-value < 0.1; n = 3-4). Genes synergistically or antagonistically regulated by the indicated recombinant cytokine pairs relative to matching single cytokines are indicated in orange and green, respectively (right). (E-H) Log2 fold-changes (E-F) or normalized counts (G-H) for indicated tissues and genes with nonlinear regulation (orange, synergistic; green, antagonistic) in mice injected with indicated cytokines (E-F) or upon LPS or CLP sepsis (G-H). Error bars, SEM (n = 3-5). **See also** Figure S3 and Table S3.

### The Pairwise Effects of TNF plus IL-18, IFN-γ, or IL-1β Lead to Organism-Wide Cellular Changes Mirroring Sepsis

The effects of the three cytokine pairs – TNF plus IL-18, IFN-γ, or IL-1β – on tissue states are likely driven by how pairwise cytokine signaling impacts the abundance and state of cell types across organs. For example, TNF plus IL-18, IFN-γ, or IL-1β led to an increase in the expression level of *S100a8* in lung, a neutrophil marker, whereas TNF plus IL-18 or IFN-γ decreased the expression of *Thy1* in thymus, a thymocyte and T cell marker (**Figure 3D and TableS3**). Building upon these observations, we sought to systematically quantify the effects of cytokine pairs and LPS on cell type abundances across the body using computational inferences from whole-tissue gene expression profiles. In brief, we first computed a cell type specificity score for each gene expressed in 195 cell types across 9 organs (*i.e.*, bone marrow, colon, heart, iLN, kidney, liver, lung, spleen, and thymus). Resulting gene-centric specificity scores were used to define ranked gene sets for each cell type. Second, we used these cell type-specific gene sets to calculate an abundance score for each cell type in each tissue from mice injected with LPS or cytokine pairs relative to untreated controls (**Figure 4A**). The cell type abundance score provides a metric to estimate the relative abundance of a given cell type in a tissue sample from mice injected with LPS or recombinant cytokines compared to matching, untreated tissue (**Methods**).

**Figure 4.**
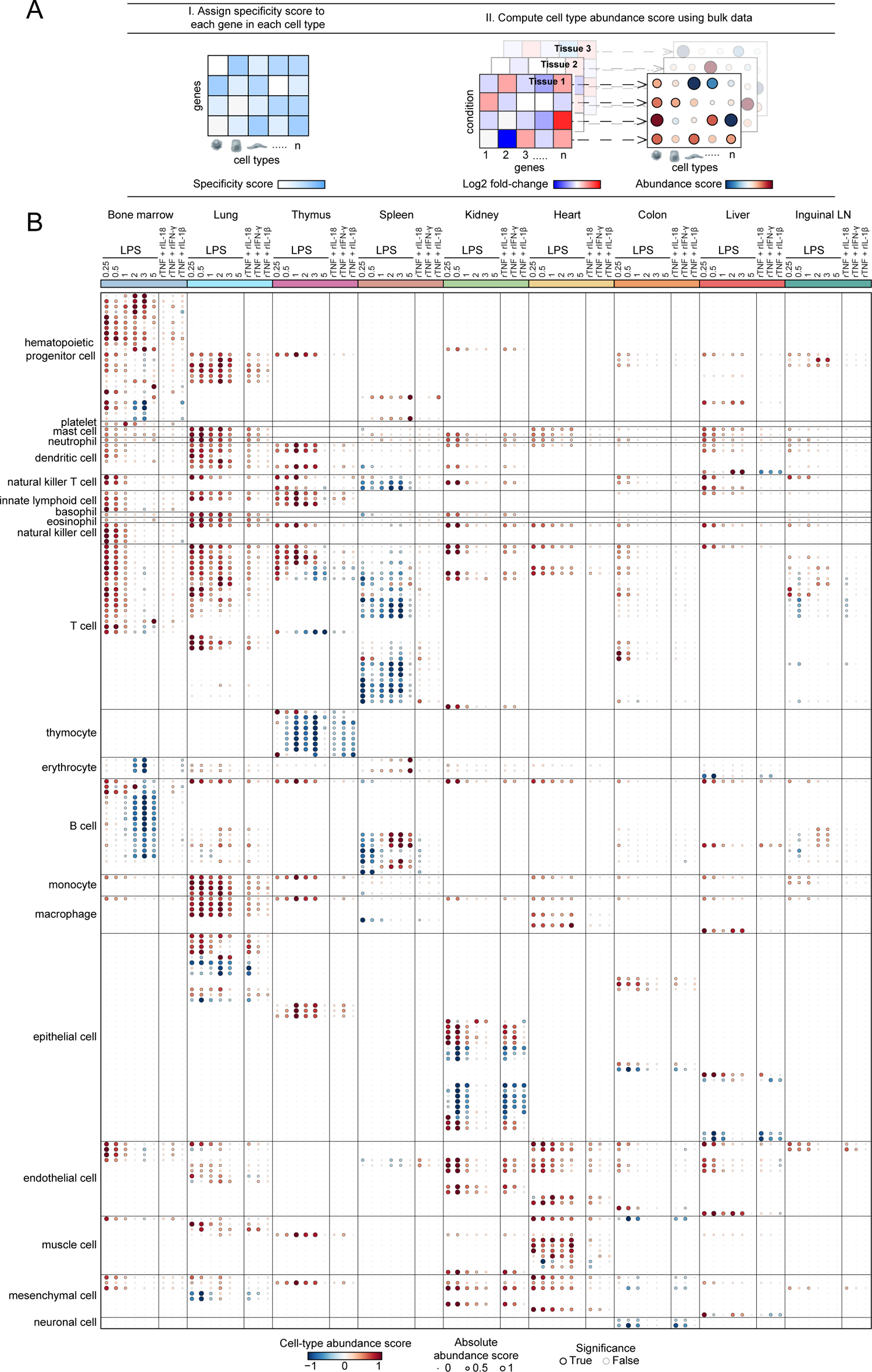
The Pairwise Effects of TNF plus IL-18, IFN-γ, or IL-1β Mirrors a Large Fraction of the Cellular Effects of Sepsis across Organs. (A) Schematic overview of the method to computationally estimate the relative abundance of cell types in organs from treated (LPS, recombinant cytokines) versus untreated, control mice by combining cell type-specific gene sets and whole-tissue gene expression measurements. (B) Cell type abundance scores computed for 195 cell types (rows) across 9 tissues (colors; top) upon injection of a sublethal dose of LPS or indicated recombinant cytokine pairs (columns). **See also** Figure S4 and Table S4.

We obtained a map of the effects of LPS and the three cytokine pairs of interest on 195 cell types, which revealed changes in cell type abundance scores across all 9 organs tested (**Figure 4B**). Notably, cytokine pairwise effects on cells mirrored those of LPS in the majority of the cell types tested and ranging from 23.3% (14/60 cellular effects by LPS at day 0.5) in bone marrow to 100% (42/42 at day 0.5) in kidney, with an average overlap in effects of 48.7% ± 27.2 SD across all 9 organs tested (**Figure S4A**). LPS and cytokine pairs led to several cellular changes which are well-described in sepsis. For example, our cell abundance analysis showed a significant decrease in B and T cell type scores across lymphoid tissues (spleen, thymus, and bone marrow) (**Figure 4B and S4B-C**), which reflects lymphopenia, a hallmark of sepsis.^9^ For T cells, all three cytokine pairs led to a strong decrease in thymocytes as in LPS, whereas the effects of these pairs on splenic T cells was less pronounced than that of LPS sepsis (**Figure 4B and S4B**). These results provide a mechanistic basis for the well-described phenomena of T cell depletion and thymic involution during sepsis.^15^ For B cells, TNF plus IL-18 led to a decrease in abundance of several splenic B cell types, whereas in the bone marrow none of the three pairs tested recapitulated LPS effects on B cells (**Figure 4B and S4C**). In another example, LPS and cytokine pairs led to an increase in abundance of endothelial cell types associated with the heart, kidney, and liver (**Figure 4B and S4D**). This observation is also corroborated by published work by others in sepsis,^38^ although the factors driving this phenotype were not known.

Next, we sought to validate experimentally several associations between cytokine pairs and changes in cell type abundances that were predicted by our computational analyses. In the bone marrow, we confirmed that TNF and IL-1β is sufficient to decrease the abundance of cell types from the erythroid lineage using flow cytometry (**Figure 5A-B**), which help to explain anemia, a well-described phenomenon in sepsis. In spleen, we found that TNF combined with IL-18 to deplete B cell subsets including follicular and, even more so, marginal zone B cells, as confirmed by the flow cytometric analysis of splenocytes (**Figure 5C-D**), in agreement with work using CLP,^39^ although the causal factors for this phenotype were not previously known. In kidney, we confirmed the prediction that all three cytokine pairs negatively impacted proximal tubule epithelial cells using spatial transcriptomics (**Figure 5E-F and S5A-E**). In addition, we used immunohistochemistry to validate the increase in neutrophils in the lungs and macrophages in the thymus in mice injected with LPS or the three cytokine pairs (**Figure 5G-H** and **S5F-G**). Lastly, in liver, hepatocytes were negatively impacted by LPS and cytokine pairs (**Figure 5I-J**). We also found that injecting LPS in *Il18^−/−^*, *Ifng^−/−^*, and *Il1b^−/−^* mice treated with anti-TNF neutralizing antibodies abrogated the cellular effects validated above in most cases, but not for lung granulocytes, suggesting that other pathways are likely at play (**Figure S5H**). Overall, by mapping the effects of LPS and cytokine pairs on cell type abundances, we provided a mechanistic basis for known and previously unreported cellular effects of sepsis on tissues (**Figure 5K**). For example, the relative abundance scores of immune cell types are positively and negatively regulated by at least one of the three cytokine pairs across all 9 organs tested (**Figure 5K**). While endothelial cell types were mostly up-regulated, epithelial and mesenchymal cell types were equally up- or down-regulated across tissues (**Figure 5K**). Out of the 3 recombinant cytokine pairs tested, TNF plus IL-18 was the one impacting the most cell types across the most tissues, reflecting its wider impact on tissue transcriptional states compared to the other 2 cytokine pairs at the doses tested (**Figure 5K**).

**Figure 5.**
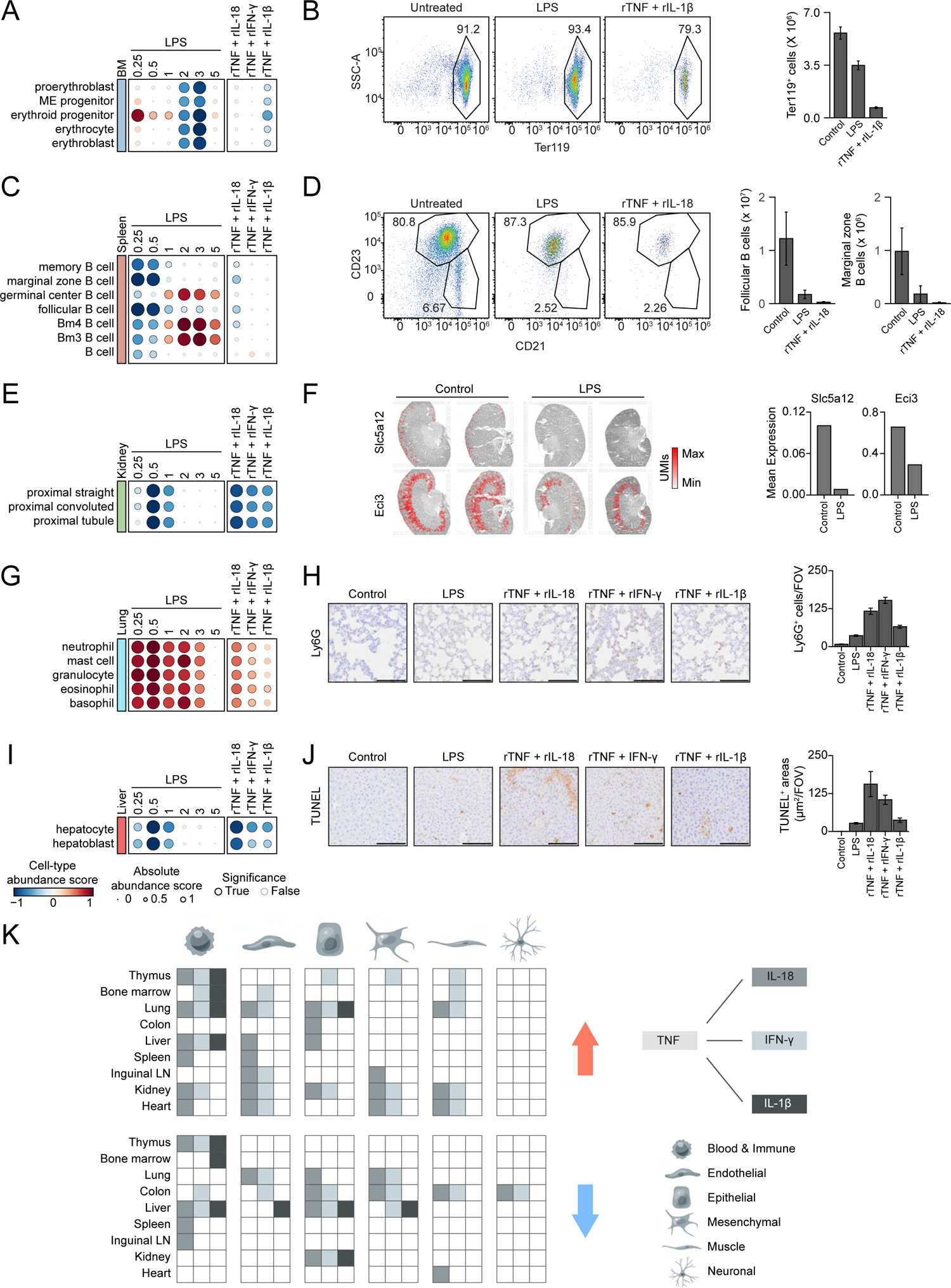
The Impacts of TNF plus IL-18, IFN-γ, or IL-1β on Cell Types Provide a Mechanistic Explanation for Known Cellular Effects of Sepsis across Tissues. (A, C, E, G, I) Cell type abundance scores computed for indicated cell types (rows) and tissues (colors) upon injection of a sublethal dose of LPS in wild-type (left) or injected with indicated recombinant cytokine pairs (right) (columns). Black borders indicate significance (z-score > 1). (B, D) Flow cytometry analysis (left) of bone marrow erythrocytes (B) or splenic B cells (D) from mice injected with a sublethal dose of LPS or indicated cytokines. Bar graphs (right) show quantifications in absolute count per tissue. Error bar, SEM (n = 2-4). (F) Spatial gene expression analysis of indicated genes (rows) from control or LPS-treated mice (columns) overlaid on grey-scale H&E images from mouse kidney sections. Bar graphs (right) show the mean expression of each gene (top) across spatial transcriptomics spots and replicate sections (n = 2). (H, J) Images (40X magnification; left) from Ly6G immunohistochemistry in lungs (H) and TUNEL staining in liver (J) from mice injected with LPS, indicated cytokines, or left untreated as controls. Bar graphs (right) show quantifications of Ly6G+ cells per field of view (FOV; H) or TUNEL^+^ areas (µm2) per FOV (J). Scale bars, 100µm; Error bar, SEM (n = 10). (K) Schematic, qualitative summary of the impact of the three cytokine pairs indicated (grey scale) on the indicated core cell types (columns and bottom-right legend) across tissues (rows). Red and blue arrows indicate an increase or decrease, respectively, in cell type abundance score for each cytokine pair on each core cell-type in each tissue. **See also** Figure S5.

### Neutralizing or Genetically Deleting PLA2G5 Promotes Survival to Sepsis

Having identified mechanisms of action for cytokines well known to play a role in sepsis, we next sought to search our organism-wide transcriptional profiles of sepsis to find previously unreported systemic regulators of sepsis. We found that phospholipase A_2_ group V (PLA2G5), a secretory and Ca2^+^-dependent lipolytic enzyme in the PLA_2_ family,^40, 41^ was strongly upregulated at the mRNA level within 6 hours of LPS injection in the colon and small intestine and downregulated in the heart and spleen (**Figure 6A-B**). Public single-cell RNA-seq data showed that *Pla2g5* is highly expressed in colon epithelial and goblet cells at steady state (**Figure 6C-D**).^42^ To test if PLA2G5 plays a role in sepsis, we injected mice with monoclonal antibodies targeting PLA2G5 prior to a challenge with LPS.^43^ We found that neutralization of PLA2G5 allowed mice to survive a lethal LPS challenge (**Figure 6E**). Similarly, *Pla2g5^−/−^* mice survived LPS or CLP sepsis better than wild-type counterparts (**Figure 6F-G** and **S6A**).

**Figure 6.**
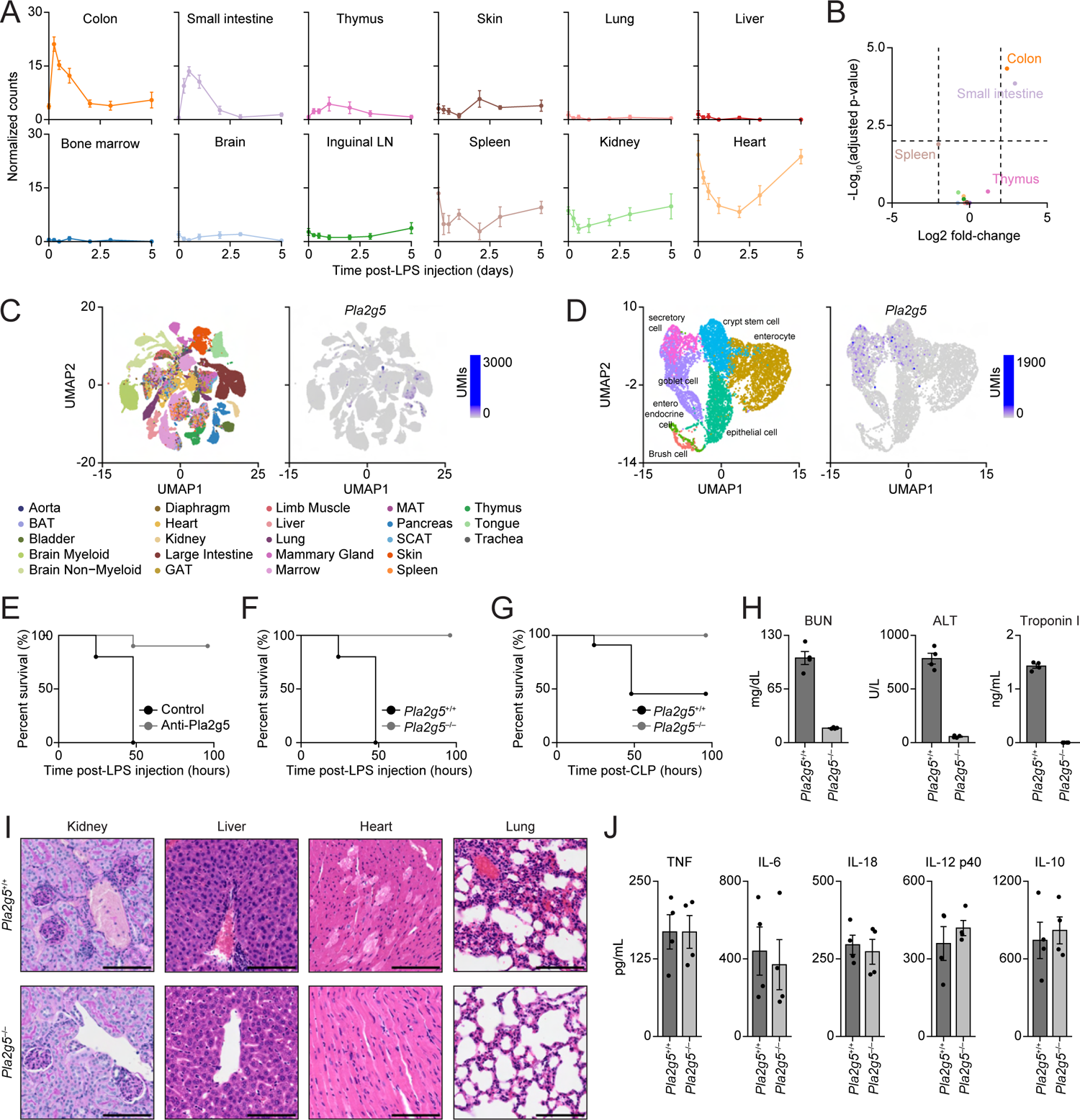
Functional Perturbations of PLA2G5 Promotes Survival to Sepsis. (A) Normalized counts for *Pla2g5* from indicated organs. LN, lymph node; Error bars, SEM (n = 3-4). (B) Volcano plot for *Pla2g5* expression at 6 hours post-LPS injection across all 12 organs from A. (C-D) Single-cell expression of *Pla2g5* mRNAs across mouse tissues. Shown are uniform manifold approximation and projection (UMAP) plots of *Pla2g5* mRNA counts (left panels; C-D) across indicated mouse organs (right panel; C) or large intestine cell types (right panel; D). BAT, brown adipose tissue; GAT, gonadal adipose tissue; MAT, mesenteric adipose tissue; SCAT, subcutaneous adipose tissue. (E-G) Survival curves of mice upon LPS (E-F) or CLP (G) sepsis using ©2G5 neutralizing antibodies (E) or indicated mouse genotypes (F-G) (n = 10-11). (H) Serum levels of indicated organ injury markers at 24 hours post-injection of a sublethal dose of LPS in indicated mouse genotypes. Error bars, SEM (n = 4). (I) Histological analysis of kidney (PAS staining) and liver, heart, and lungs (H&E staining) from *Pla2g5*^+/+^ (top) or *Pla2g5*^-/-^ (bottom) mice at 24 hours post-sublethal LPS injection. (J) Serum levels of indicated cytokines (top) in *Pla2g5*^+/+^ or *Pla2g5*^-/-^ mice at 24 hours after LPS injection. Error bar, SEM (n = 4). **See also** Figure S6 and Table S5.

To investigate the mechanism of action of PLA2G5 in sepsis, we first measured tissue injury markers and found that both PLA2G5 neutralization and genetic deletion led to decreased plasma levels of renal, liver, and cardiac injury markers – blood urea nitrogen (BUN), alanine aminotransferase (ALT), and troponin I, respectively – during sepsis (**Figure 6H** and **S6B**). Furthermore, tissue sections from LPS-injected mice showed decreased signs of histopathology in *Pla2g5^−/−^* mice compared to wild-type controls (**Figure 6I**). Interestingly, we found that the blood levels of inflammatory cytokines were unchanged in *Pla2g5^−/−^* compared to wild-type mice upon LPS injection (**Figure 6J**), which suggested that the negative effects of PLA2G5 on the host during sepsis was independent of cytokines.

Next, we hypothesized that PLA2G5 could generate lipid mediators responsible for the negative impact of this phospholipase on the host during sepsis. We found that PLA2G5 blockade reduced the levels of several lipid metabolites in plasma, but not colon, including those of fatty acids (FA) such as oleic acid (18:1) and linoleic acid (18:2), lysophospholipids such as lysophosphatidic acid (LPA), lysophosphatidylcholine (LPC), lysophosphatidylethanolamine (LPE), and lysophosphatidylserine (LPS) species, and metabolites derived from polyunsaturated fatty acids (PUFAs) such as arachidonic acid (AA), eicosapentaennoic acid (EPA), docosahexaenoic acid (DHA), and linoleic acids (**Figure S6C-G**). We found that 6-ketoprostaglandin F_1α_ (6-keto-PGF_1α_), a stable metabolite derived from PGI_2_ and resolving D2 (RvD2) were downregulated in colon from LPS-treated mice upon PLA2G5 blockade (**Figure S6E**). The levels of all other lipid species measured in the plasma and colon were relatively low or showed little to no differences upon PLA2G5 blockade during LPS sepsis (**Figure S6C-G**). Using these lipidomics profiles and prior knowledge on lipid and PLA_2_ enzyme biology,^40, 41^ we selected the following 12 lipid species for further study: LPA 16:0, LPA 18:0, LPA 18:1, 12-hydroxyeicosatetraenoic acid (12-HETE), 13-hydroxyoctadecadienoic acid (13-HODE), 9-hydroxyoctadecadienoic acid (9-HODE), 9-oxo-octadecadienoic acid (9-oxo-ODE), beraprost (a stable analogue of PGI_2_ because 6-keto-PGF_1a_, which was measured by lipidomics, is an inactive metabolite derived from PGI_2_), oleic acid (18:1), linoleic acid (18:2), LPC, and LPE. In addition, we selected two inhibitors of LPA signaling: PF8380, which blocks the autotaxin enzyme producing LPA from LPC and LPE, and Ki6425, which antagonizes LPA_1_ or LPA_3_ receptors.

To test if these lipids could negatively impact sepsis outcomes, as PLA2G5 does, we injected mice with purified lipids and challenged these animals with sublethal or lethal doses of LPS. Out of the 12 lipids and 2 inhibitors tested, LPA 18:0 was the only metabolite which fully rescued mice from both a lethal and a sublethal LPS challenge (**Figure S6H-I**), in agreement with a previous report.^44^ Beraprost and 13-HODE led to a partial rescue of the drop in body temperature triggered by a sublethal LPS challenge (**Figure S6I**). We found that LPA 18:0 injection maintained the blood levels of tissue injury markers at homeostasis during sepsis (**Figure S6J**). Furthermore, we observed that LPA 18:0 triggered an organism-wide anti-inflammatory state reminiscent of the drug dexamethasone (**Figure S6K-M**), a corticosteroid with known benefits in viral sepsis and some cytokine storm and inflammatory syndromes.^13, 45, 46^ Taken together, these results provided insights about how LPA 18:0 exerts strong anti-inflammatory effects on the host during sepsis, but we still lacked an explanation as to how PLA2G5 blockade and genetic deletion fully protected the host from a lethal sepsis challenge.

### Blood PLA2G5 Increases Hemolysis and Plasma Heme Levels, Decreasing Survival to Sepsis

To decipher the negative impact of PLA2G5 on the host during sepsis, we measured changes in whole-tissue gene expression during sepsis across 12 organs (bone marrow, brain, colon, heart, inguinal lymph node, kidney liver, lung, skin, small intestine, spleen, and thymus) from mice treated with anti-PLA2G5 neutralizing antibodies (**Table S6**). PLA2G5 blockade impacted a relatively low number of genes regulated during sepsis: 8.7% (386/4,443) of all genes regulated at day 0.5 post-LPS across 7 out of the 12 organs profiled (**Figure 7A**), and these effects on tissue states were most pronounced in the spleen, bone marrow, and lungs (**Figure 7A-B**). Interestingly, we observed a link between PLA2G5 blockade and changes in pathways related to splenic macrophages and iron homeostasis (**Figure 7A**). Others have shown that splenic red pulp macrophages play a role in red blood cell recycling and iron homeostasis,^47, 48^ prompting us to explore the potential effects of PLA2G5 on these cells and processes.

**Figure 7.**
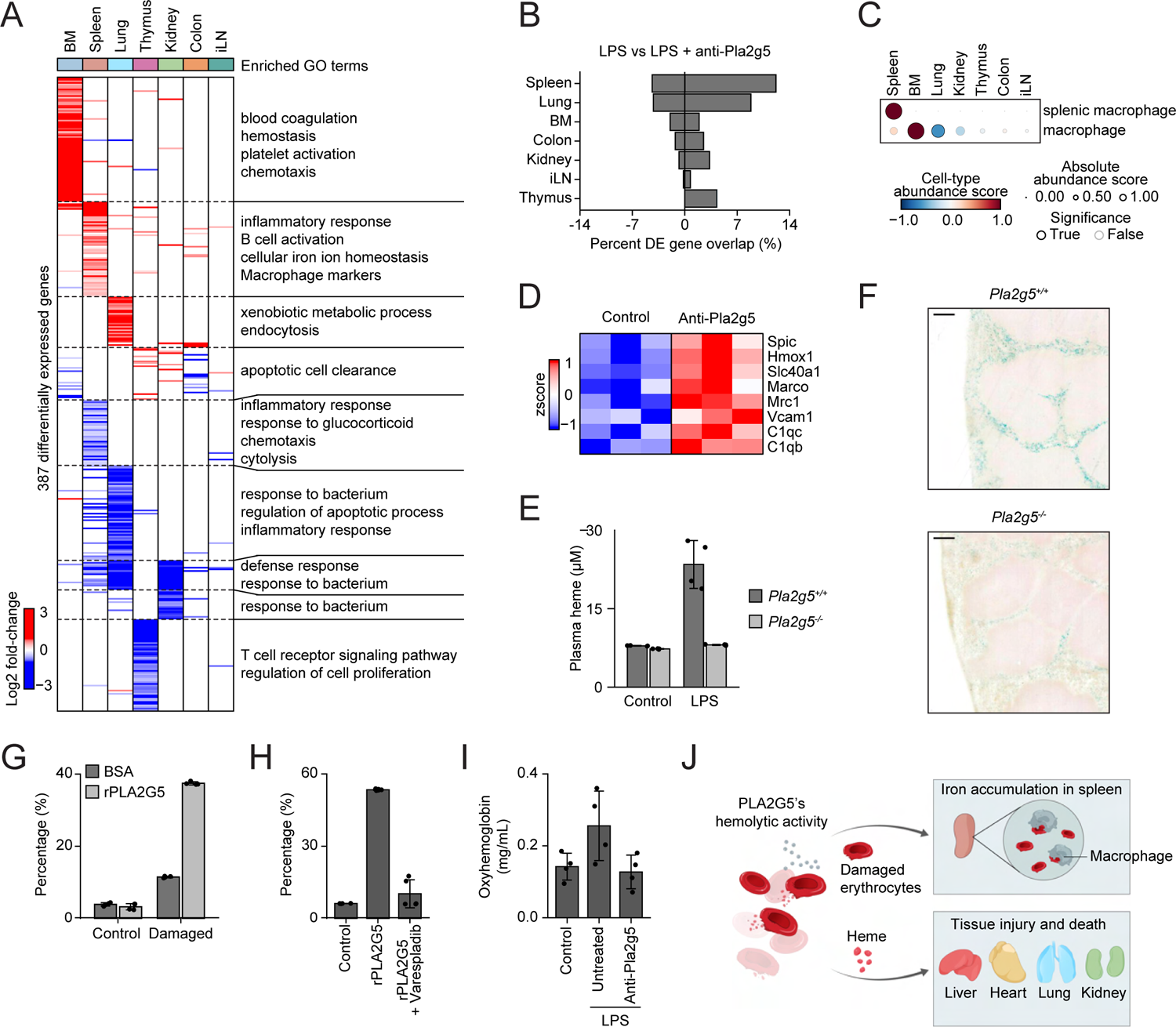
Bloodborne PLA2G5 Mediates Hemolysis and Increases Plasma Heme Levels. (A) Heatmap of differentially expressed genes (rows) from whole-tissue mRNA profiles ordered by k-means clustering (dashed lines) and organ types (top, colors) at 12 hours after sublethal LPS injection and anti-PLA2G5 antibody pre-treatment. Values are log2 fold-changes relative to matching tissues from LPS-treated mice without antibody blockade (FDR-adjusted p-value < 0.01; n = 3-4). Representative enriched Gene Ontology (GO) terms within each cluster are indicated on the right (p-value < 0.05). (B) Percentages (x axis) of genes differentially expressed in tissues (rows) upon pre-treatment with anti-PLA2G5 neutralizing antibodies in LPS-injected mice that match the genes regulated by LPS alone. BM, bone m©ow; iLN, inguinal lymph node. (C) Cell type abundance scores computed for indicated cell types (rows) across tissues (columns) upon injection of a sublethal dose of LPS in mice pre-treated with anti-PLA2G5 neutralizing antibodies relative to untreated mice. Black borders indicate significance (z-score > 1). (D) Heatmap of gene expression levels for genes related to splenic red pulp macrophage (rows) from whole-spleen mRNA profiles at 12 hours after sublethal LPS injection with or without anti-PLA2G5 antibody pre-treatment (columns). Values are normalized counts scaled by row (FDR-adjusted p-value < 0.01; n = 3). (E) Plasma heme concentrations in *Pla2g5*^+/+^ or *Pla2g5*^-/-^ mice 24 hours after LPS or PBS (control) injections. Error bar, SD (n = 4). (F) Images (40X magnification) of Prussian blue staining for ferric iron on spleen histological sections from LPS-injected *Pla2g5*^+/+^ (top) or *Pla2g5*^-/-^ (bottom) mice. Scale bar, 200 µm. (G-H) *In vitro* hemolysis assay. Shown are percentages of cell-free oxyhemoglobin (relative to an equal volume of chemically lysed erythrocytes) in cell lysates from normal (Control) or phosphatidylserine (PS)-exposing (Damaged) erythrocytes incubated with recombinant PLA2G5 (rPLA2G5) or BSA for 1 hour at 37°C (G), or PS-exposing erythrocytes incubated with rPLA2G5 in the presence or absence of the secreted phospholipase A2 inhibitor Varespladib or left untreated (control) (H). Error bars, SD (n = 4). (I) Plasma oxyhemoglobin levels in mice injected with plasma which were collected from LPS- or PBS (Control)-injected mice and incubated *in vitro* with anti-PLA2G5 neutralizing antibodies for 4 hours at 4°C prior to transfer into naive mice. Error bars, SD (n = 4). (J) Schematic depicting the proposed model for the role of PLA2G5 in sepsis. **See also** Figure S7 and Table S6.

Using our computational scoring method to estimate cell type abundances from whole-tissue expression profiles (**Methods**), we found that splenic macrophages were more abundant upon PLA2G5 blockade than in untreated mice during LPS sepsis (**Figure 7C**). Several genes linked to splenic red pulp macrophages, such as *Spic*, *Hmox1*, and *Vcam1*,^47, 48^ were upregulated upon PLA2G5 blockade in LPS-injected mice (**Figure 7D**), suggesting that PLA2G5 impacts the abundance of these splenic cells. Next, we measured the blood levels of heme in *Pla2g5^+/+^* and *Pla2g5^−/−^* mice challenged with LPS and found a significant decrease in heme in knockout animals (**Figure 7E**). The decrease in heme blood levels in *Pla2g5^−/−^* mice upon LPS challenge compared to wild-type controls was associated with a decrease in Prussian blue staining of ferric iron in the spleen (**Figure 7F** and **S7A**). Based on these observations, we asked if PLA2G5 could directly lyse red blood cells (RBCs) using an *in vitro* assay whereby recombinant PLA2G5 protein was mixed with mouse RBCs. Recombinant PLA2G5 led to increased RBC lysis *in vitro* upon treatment with Ca^2+^ and a calcium ionophore, which damages the plasma membrane of RBCs by disrupting the distribution of phospholipids (**Figure 7G**). Furthermore, the hemolytic effects of PLA2G5 was dependent on its phospholipase activity as shown by treatment with varespladib, a broad inhibitor of secreted PLA2s (**Figure 7H**).^49^

Lastly, to assess whether PLA2G5 was present in the blood circulation of mice undergoing sepsis and led to RBC lysis *in vivo*, we performed plasma transfer experiments. After collecting plasma from wild-type and *Pla2g5^−/−^* mice treated with LPS, we incubated plasma with anti-PLA2G5 or isotype control antibodies, and injected those plasma© naive, wild-type mice. Plasma from LPS-injected, wild-type mice led an increase in RB©is in naive mice, whereas plasma from PLA2G5 knockout or blockage did not (**Figure 7I** and **S7B**). Overall, these results support a model whereby the release of PLA2G5 into the systemic circulation during sepsis leads to a concurrent increase in hemolytic activity in the blood, hemophagocytosis in the spleen, and heme levels in the blood, the latter being linked to multi-tissue injury (**Figure 7J**).^50, 51^

## DISCUSSION

While sepsis remains a leading cause of death in intensive care units worldwide, our understanding of the pathogenesis of sepsis across most tissues and organs of the body remains rudimentary. To begin to address this fundamental gap in knowledge, we studied the organismal response to sepsis over time by measuring changes in gene expression across tissues in mouse models of the disease. Building upon our spatiotemporal map of sepsis, we discovered two interorgan mechanisms which help explain the pathophysiology of the disease through the uncontrolled activities of well-described and previously unreported factors: a hierarchical cytokine module composed of TNF, IL-18, IFN-γ, and IL-1β, and the secreted phospholipase PLA2G5, respectively.

What did organism-wide maps of gene expression tell us about sepsis? First, our data revealed a plethora of changes that come with the initiation and resolution of sepsis, both at the molecular and cellular levels. These changes were detected in all organ systems tested and encompassed most known, if not all, biomarkers and physiological events linked to sepsis. For example, out of the 809 genes with a PubMed Gene Reference19enerifyunction (geneRIF) annotation containing the keyword “sepsis”, 70% (563/809 genes) were regulated in at least one tissue and time point during sepsis. Future work is needed to elucidate which regulated genes are causal or bystander and beneficial or detrimental during sepsis.

Second, we found that non-lymphoid tissues regained homeostasis sooner than lymphoid ones. This result is reminiscent of how some organs reverse dysfunction in sepsis, including those poor at regenerating such as heart, lung, kidney, or brain, whereas the immune system suffers long-term dysregulation with life-threatening consequences for survivors.^20, 32^ Further mining our data might help to identify factors that safeguard non-lymphoid tissues, such as IL-10 for microglia^52^ or GDF15 for heart^53^ – both present in our data – or those that damage lymphoid tissues and cells. Third, we built an organism-wide map of sepsis effects at the resolution of cell types by computing abundance scores for 195 cell types across 9 organ types. In addition to revealing the scope of the cellular effects of sepsis on tissues, our analysis provided a linkage between specific cytokine pairs and cell types across tissue contexts. Notably, all six associations between cytokines and cell types selected for further study were validated by experiments (**Figure 5 and S5**). Most of these cellular effects were previously observed in sepsis, such as an increase in thymic macrophages,^54^ erythropenia,^27^ splenic B cell loss,^15^ or changes in kidney tubules^55^ but lacked causal factors. Future work is needed to test other predictions and define the mechanisms underlying cellular changes, such as alterations in the proliferation, death, migration, or intracellular state of the cells affected by sepsis. Taken together, our spatiotemporal data provide detailed insights in the quest towards defining a mechanistic framework to explain sepsis.

What sense can be made of the cytokine cacophony taking place in the blood during sepsis? While uncontrolled cytokine signaling is harmful to the body,^13^ we lack a mechanistic framework to explain which signaling events impact each cell type in the context of each tissue. Several features of the cytokine language make it hard to decode, such as the variations in concentrations (local and systemic), activities (pro, anti, or both for any given cytokine), and interactions within a mixture of cytokines present in a tissue. Here, by measuring the impact of six cytokines alone or in pairwise combinations on tissue mRNA expression profiles, we captured the net output of tissue-level responses to cytokine inputs – as opposed to focusing on the response of a single cell type. Our data support a model whereby a few elements of cytokinic information – TNF plus IL-18, IFN-γ, or IL-1β – suffice to explain a large fraction of the molecular and cellular effects of sepsis across tissues.

Our discovery of a simplifying hierarchy among the cytokines upregulated in blood during sepsis will help to build a unifying mechanistic framework for sepsis and other cytokine storms.^13^ Notably, the proposed cytokine hierarchy relies on nonlinear interactions between TNF and IL-18, IFN-γ or IL-1β signaling, a notion well supported by four decades of work on cytokine interactions *in vitro* and *in vivo*. For example, TNF has been shown to combine synergistically or antagonistically with IFN-γ or IL-1β to impact secretion, cell death or proliferation, and cell states in immune and non-immune cells in culture.^56–65^ While the interaction between TNF and IL-18 had not been reported to our knowledge, TNF plus IFN-γ^66–69^ or IL-1β^70–72^ worsen the outcome of sepsis and other inflammatory disorders *in vivo*. The cytokines of this module also influence each other’s production,^13^ which further supports the hierarchy uncovered by our pairwise cytokine screening data.

Future investigations are needed to define the direct and indirect effects of each cytokine pair on each cell types. For example, it is likely that some cytokine pairwise effects act through downstream factors, including through the release of other cytokine or non-cytokine diffusible factors that are directly sensed by the cells and tissues. In addition, while our data linked one of the three cytokine pairs or more with 52% (178/342 at day 0.5) of the target cell types tested in at least one organ type and impacted with LPS, the other half of the cellular effects of LPS on tissues remained unexplained by the three cytokine pairs used here. Thus, further work on other cytokines and non-cytokine factors, such as the complement or coagulation systems, is needed to pinpoint the causative factors responsible for the observed cellular effects of sepsis on tissues. The detailed signaling events mediating cytokine interactions at the level of cells also remain to be elucidated, such as the putative rewiring of the MAPK, NF-κB, IRF and Jak/STAT pathways that have previously been linked to the interaction between TNF and IFN-γ.^65, 67, 73, 74^

Why is TNF the central node of this cytokine module recapitulating many of the effects of sepsis? After half a century since the first isolation of TNF as a factor which could kill tumor cells,^75^ TNF has been implicated in the pathogenesis of countless infectious and non-infectious diseases.^76, 77^ In sepsis, TNF is one of the earliest cytokines produced in mice and humans, peaking in the blood in less than 2 h with a circulating half-life of less than 20 min of mice.^78, 79^ Anti-TNF antibodies protect against lethal sepsis when present before or early on upon the start of the disease,^80, 81^ but not later in the disease, which helps to explain the failure of anti-TNF therapy in humans with sepsis.^3, 82–85^ Interestingly, pre-treatment with anti-TNF antibodies leads to beneficial effects in humans, such as in the suppression of the Jarisch-Herxheimer reaction occurring in response to antibiotic treatment of louse-borne relapsing fever.^86, 87^ Conversely, the infusion of recombinant TNF in humans suffices to trigger flu-like symptoms.^88^ Lastly, our findings about the central role of TNF in a cytokine circuit controlling sepsis are reminiscent of the existence of a cytokine hierarchy defining human chronic inflammatory diseases across tissues.^89, 90^ Inhibiting TNF has shown remarkable therapeutic benefits in patients with psoriasis, psoriatic arthritis, Crohn’s disease, ulcerative colitis, ankylosing spondylitis, juvenile arthritis and many other less prevalent diseases. However, targeting cytokines such as IL-6, IL-1, or IL-17/23 showed a much narrower range of efficacy, suggesting that TNF combines with select cytokines in select organs in the pathogenesis of inflammatory disorders.^89, 90^ The multi-tissue effects of TNF in inflammation are likely a product of the existence of numerous TNF receptors which are ubiquitously expressed.^77^ Taken together, these observations in sepsis and beyond help to contextualize the organism-wide effects of the three TNF-centered cytokine pairs identified by our data as critical to explain sepsis.

Beyond cytokine signaling, we uncovered an interorgan axis whereby the release of PLA2G5 into the blood circulation leads to hemolysis, increased heme levels, and organ injury. PLA2G5 has several local pathogenic and protective roles linked to various cell types, including macrophages, adipocytes, endothelial cells, bronchial epithelial cells, and cardiomyocytes,^40, 41^ but its role in sepsis was previously unknown. Interestingly, another member of the secreted PLA2 family, namely PLA2G2A and often referred to as sPLA2-IIA, is a sepsis biomarker which can release arachidonic acid, the precursor of the eicosanoid biosynthetic pathways yielding pro- and anti-inflammatory lipid species.^40, 91, 92^ In addition, PLA2G2A has been reported to hydrolyze phospholipids *in vitro* in erythrocyte-derived microvesicules^93^ or activated or damaged erythrocyte membranes directly,^94^ although the ability of PLA2G5 to hydrolyze cell membranes has been shown to be greater than that of PLA2G2A.^95^ PLA2G5 is also able to act on the membranes of endothelial cells and bacteria.^40, 41, 96^ Perhaps the activity of PLA2G5 on microbial membranes is indicative of a putative physiological function for PLA2G5 in the gut, where its gene is highly expressed in epithelial cells and which might impact the microbiota similar to PLA2G2A.^97^ Some of the lipid metabolites impacted by PLA2G5 in the blood of septic mice, including fatty acids or lysophospholipids, are likely due to additional, indirect effects of PLA2G5 because these lipids are not direct targets of PLA2G5’s lipolytic activities.^40^ The hemolytic activity of PLA2G5 in sepsis is also reminiscent of the toxicity of sPLA_2_ present in snake venom.^98, 99^ The increase in heme blood levels due to the hemolytic activity of PLA2G5 in septic blood and downstream multi-tissue injury is consistent with previous work on the toxicity of heme on tissues during sepsis.^50, 51^ Future work is needed to clarify if PLA2G5 exerts any other role in sepsis, and to decipher the mechanism of action of PLA2G5-derived LPA 18:0, which rescued mice from sepsis through potent anti-inflammatory properties across tissues. Taken together, our data suggest that sepsis corrupts PLA2G5 – perhaps through the disruption of gut epithelia^100, 101^ – into becoming an “intestinal venom” for the host.

## AUTHOR CONTRIBUTIONS

Conceptualization and Methodology, M.T. and N.C.; Investigation, M.T., M.P., D.C., G.R., S.P., K.C., T.U., and N.C.; Lipidomics, Y.M., Y.T., and M.M.; Cell type specificity analysis, A.P., M.T., and N.C.; Topic modeling analysis, P.C. and M.S.; Gene expression interaction scoring, K.J., M.S., and N.C.; Formal Analysis, M.T., A.G., and N.C.; Resources, S.M.D., and N.C.; Writing – Original Draft, M.T. and N.C.; Writing – Review & Editing, All Authors; Funding Acquisition, N.C.; Supervision, N.C.

## ACKNOWLEDGEMENTS

We thank members of the Chevrier lab for valuable discussions; UChicago core facilities: the Single Cell Immunophenotyping Core (Mustafa Abasiyanik, Ha-Na Shim), the Animal Resources Center (Ani Solanki), the Integrated Light Microscopy Core, and Human Tissue Resource Center; the In Vivo Animal Core at the University of Michigan; staff at the Research Computing Center at UChicago for providing high-performance computing resources; and SciStories for help with artwork. M.T. was supported by the Astellas Foundation for Research on Metabolic Disorders and K.J. by the National Science Foundation Graduate Research Fellowship under Grant No. 2140001. M.M. was supported by grants AMED-CREST JP22gm1210013 from the Japan Agency for Medical Research and Development and Grants-in-Aid for Scientific Research (S) JP20H05691 from the Japan Society for the Promotion of Science. N.C was supported by NIH grants DP2-AI145100 and U01-AI160418, the Chan-Zuckerberg Initiative, the University of Chicago Medicine Comprehensive Cancer Center (UCCCC) Janet D. Rowley Discovery Fund, the University of Chicago Center for Interdisciplinary Study of Inflammatory Intestinal Disorders (C-IID) (P30 DK42086), and funds from the Chicago Immunoengineering Innovation Center and the Pritzker School of Molecular Engineering at the University of Chicago.

## DECLARATION OF INTERESTS

A.P. is the founder and CEO of Combinatics Inc. All other authors declare no competing financial interests.

## SUPPLEMENTARY FIGURE TITLES AND LEGENDS

**Figure S1.**
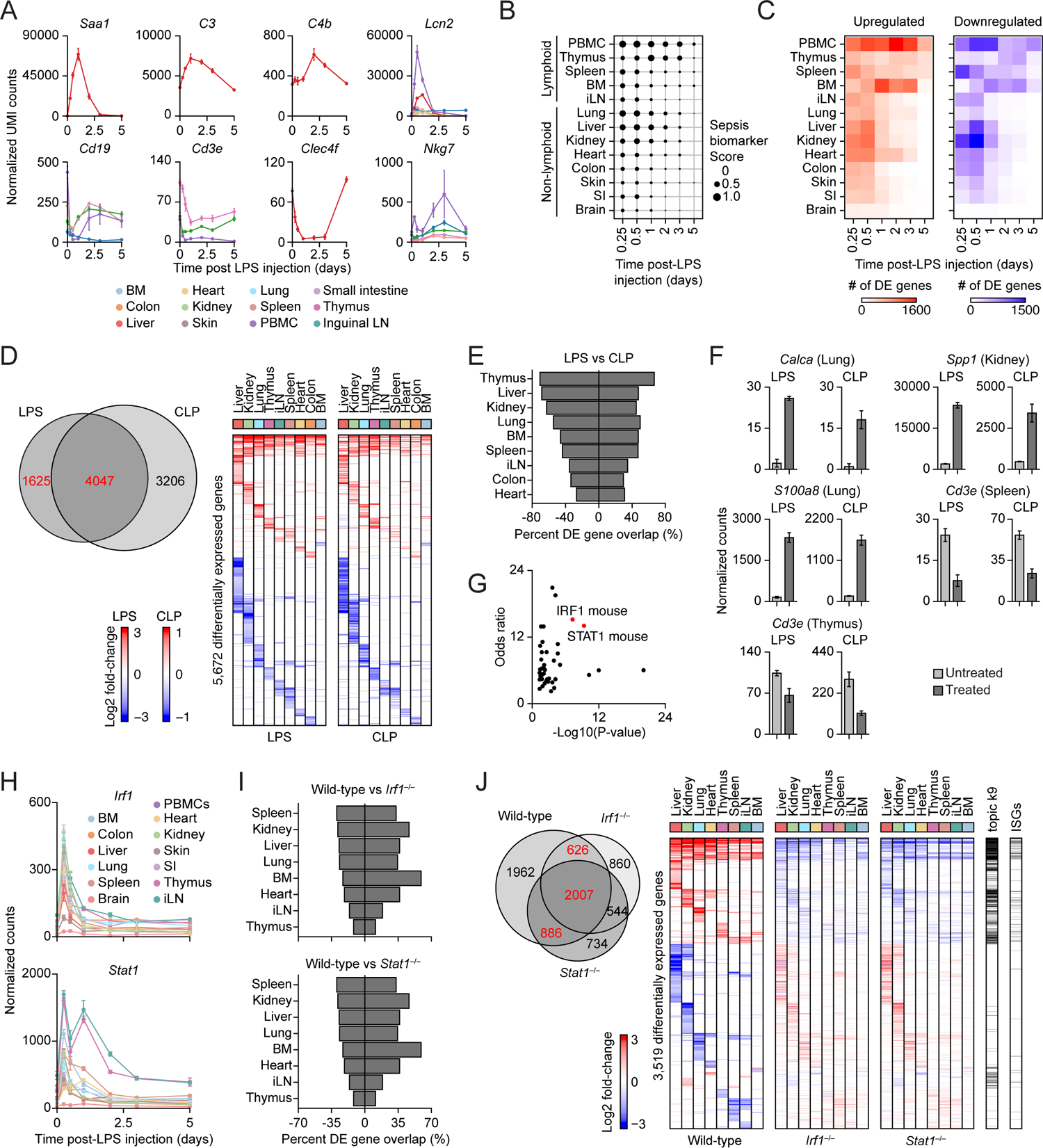
Comparative, Multi-Tissue Expression Analysis across Bacterial Sepsis Models and Transcription Factor-Deficient Mice, Related to Figure 1. (A) Normalized counts for indicated genes and organs (color). BM, bone marrow; iLN, inguinal lymph node; PBMCs, peripheral blood mononuclear cells; Error bars, SEM (n = 2-4). (B) Expression of sepsis biomarker genes (score) in each organ (rows) at each time point post-sublethal LPS injection (columns). Rows are ordered from top to bottom by high to low scores and by lymphoid and non-lymphoid tissues. SI, small intestine. (C) Heatmap showing the total numbers of up-(left) and down-regulated (right) genes across time post-LPS (columns) for each organ (rows). (D) Heatmap of differentially expressed genes (rows) from whole-tissue mRNA profiles ordered by k-means clustering and organ types (top, colors) at 12 hours after sublethal LPS injection (left) or 16 hours after severe cecal ligation and puncture (CLP; right) (Methods). Values are log2 fold-changes relative to matching organs from untreated mice for LPS and after sham surgeries for CLP (FDR-adjusted p-value < 0.1; n = 5). The Venn diagram (top right) indicates the overlap in differentially expressed genes between LPS and CLP across all 9 organs measured (red, genes shown in heatmaps). (E) Percentages (x axis) of genes differentially expressed in tissues (rows) upon severe CLP that match the genes regulated by sublethal LPS at 12 hours post-sepsis induction. (F) Normalized counts for indicated genes and organs in LPS and CLP sepsis. Error bars, SEM (n = 4-5). (G) Transcription factor enrichment analysis (Methods) on genes from cluster k = 9 of the topic modeling analysis shown in Figure 1E-F. (H) Normalized counts for indicated genes and organs (color). Error bars, SEM (n = 2-4). (I) Percentage (x axis) of genes differentially expressed in tissues (rows) at 16 hours post-sublethal LPS injection from *Irf1^−/−^* (top) and *Stat1^−/−^* (bottom) mice that match the genes regulated by LPS in wild-type mice. (J) Heatmap of differentially expressed genes (rows) from whole-tissue mRNA profiles ordered by k-means clustering and organ types (top, colors) at 12 hours after sublethal LPS injection in wild-type (left), and 16 hours after sublethal LPS injection in *Irf1^−/−^* (center) and *Stat1^−/−^* (right) mice. Values are log2 fold-changes relative to matching organs from untreated controls for wild-type (FDR-adjusted p-value < 0.01, absolute fold change > 2; n = 4) and from LPS-injected wild-type for knockouts (FDR-adjusted p-value < 0.05, absolute fold change > 2; n = 2-4). The Venn diagram (top right) indicates the overlap in differentially expressed genes between all three genotypes across all 8 organs measured (red, genes shown in heatmaps). Genes from topic k9 (topic modeling in Figure 1) and interferon-stimulated genes (ISGs) are indicated in black on the right of the heatmaps.

**Figure S2.**
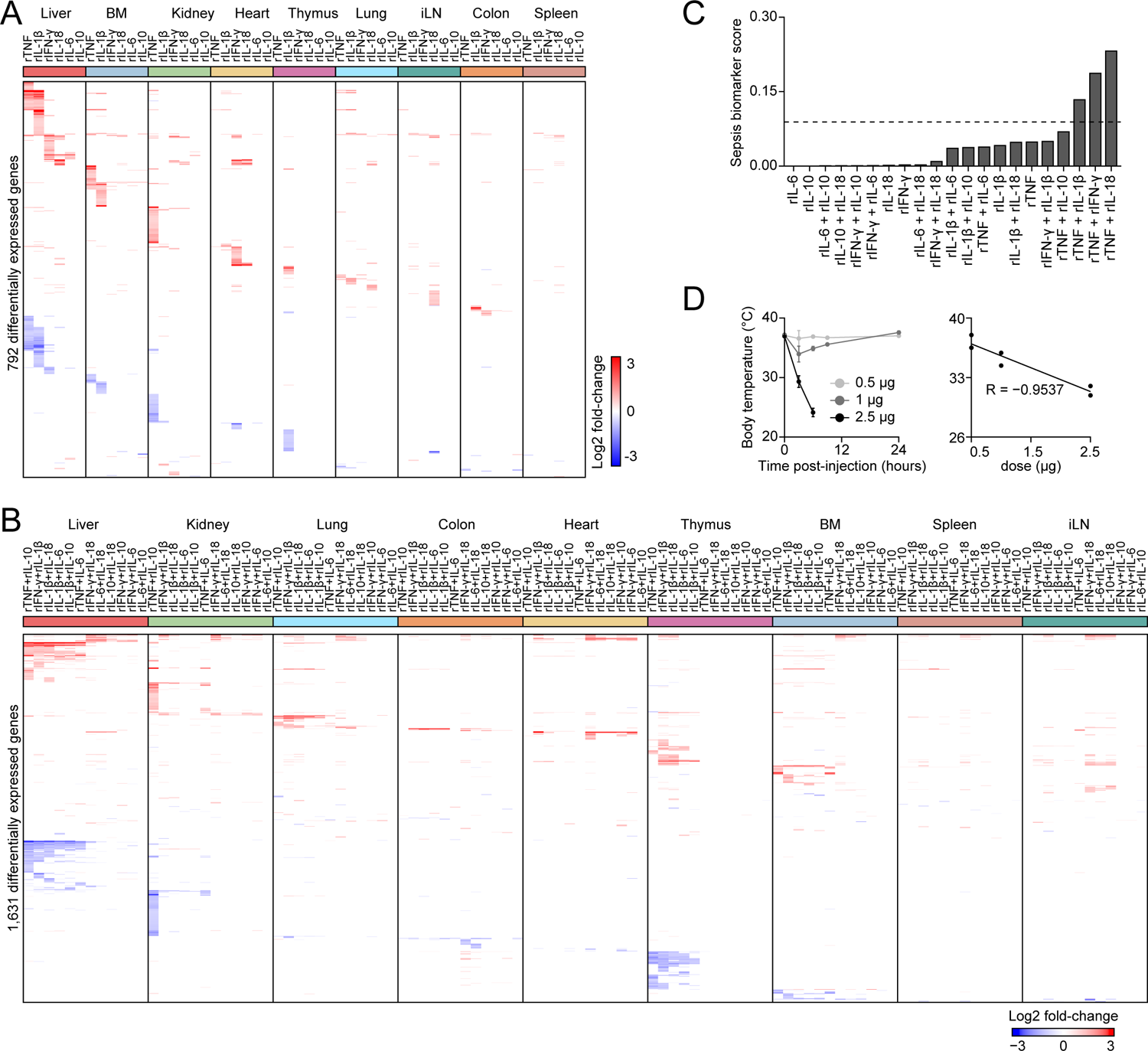
Recombinant Cytokines Injected Alone or in Pairwise Combinations Impact Tissue Transcriptional States, Related to Figure 2. (A-B) Heatmaps of differentially expressed genes (rows) from whole-tissue mRNA profiles ordered by k-means clustering and organ types (top, colors) at 16 hours after injection with indicated recombinant cytokines used alone (A) or in pairwise combinations (B). Values are log2 fold-changes relative to matching organs from untreated, control mice (FDR-adjusted p-value < 0.05, absolute fold change > 2; n = 3-4). (C) Sepsis biomarker score average (scaled by condition) across all 9 organs profiled in A and B in indicated recombinant cytokine conditions (x axis). Dashed line indicates one standard deviation. Sepsis biomarker score is the average, absolute log2 fold-change values for all 258 genes identified as sepsis biomarker in the literature. (D) Measurements of rectal temperature (y axis; left and right) relative to time post-injection (left) or the dose (right) of TNF + IL-1β. Error bars, SD (n = 2).

**Figure S3.**
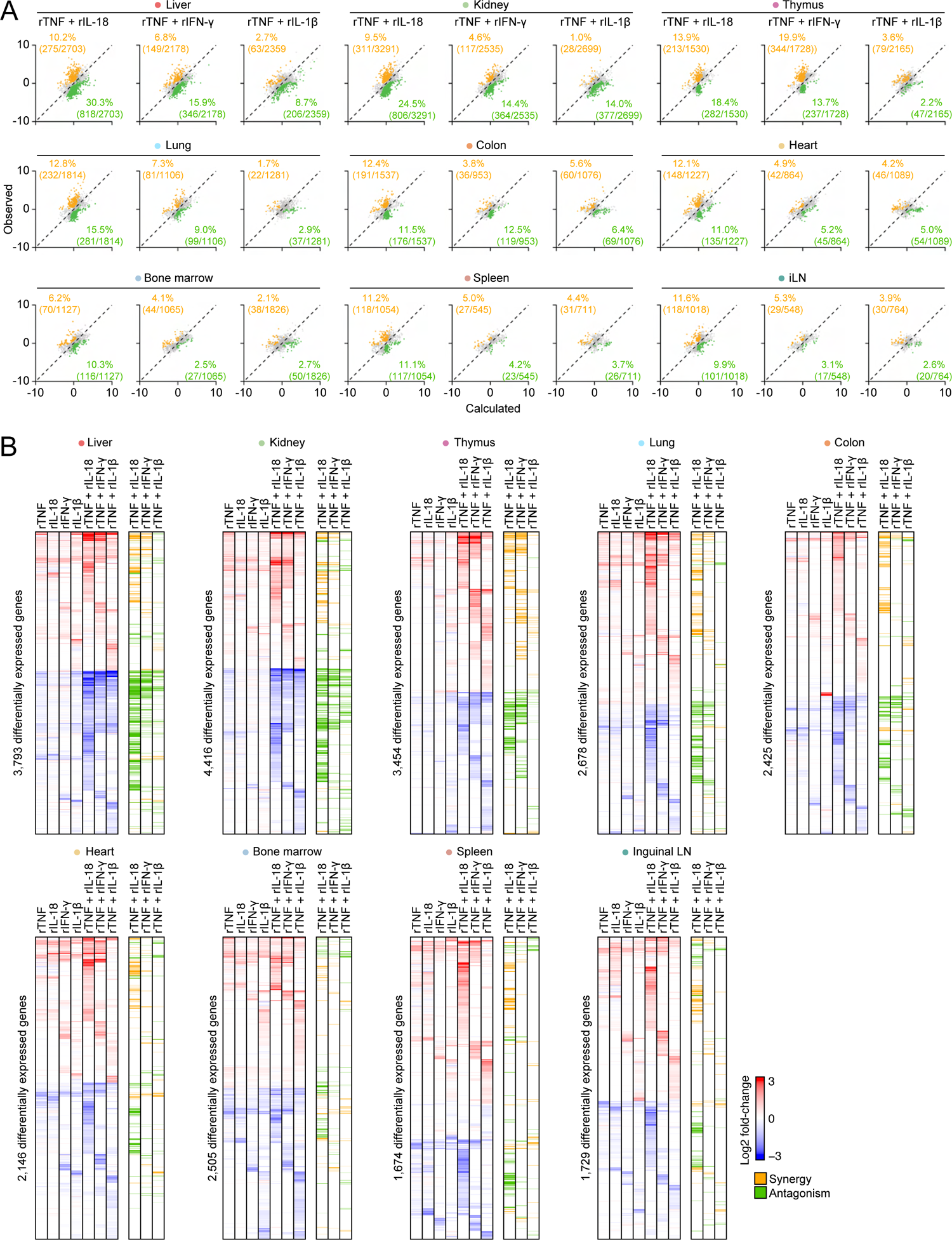
The Cytokine Pairs Composed of TNF plus IL-18, IFN-γ, or IL-1β Yield Nonlinear Interactions at the Gene Expression Level across Organs, Related to Figure 3. (A) Dot plots of the observed (y axis) and calculated (x axis) pairwise cytokine interaction effects relative to matching single cytokines on differentially expressed (DE) genes (dots) in indicated organs (top). Percentages and absolute counts of DE genes classified as synergistic (orange) or antagonistic (green) upon pairwise cytokine injection relative to matching singles. (B) Heatmaps of differentially expressed genes (rows) from whole-tissue mRNA profiles for each indicated organ ordered by k-means clustering at 16 hours after injection of indicated recombinant cytokines. Values are log2 fold-changes relative to matching, untreated organs (FDR-adjusted p-value < 0.05; absolute fold change > 2; n = 3-4). Genes synergistically or antagonistically regulated by the indicated recombinant cytokine pairs relative to matching single cytokines are indicated in orange and green, respectively.

**Figure S4.**
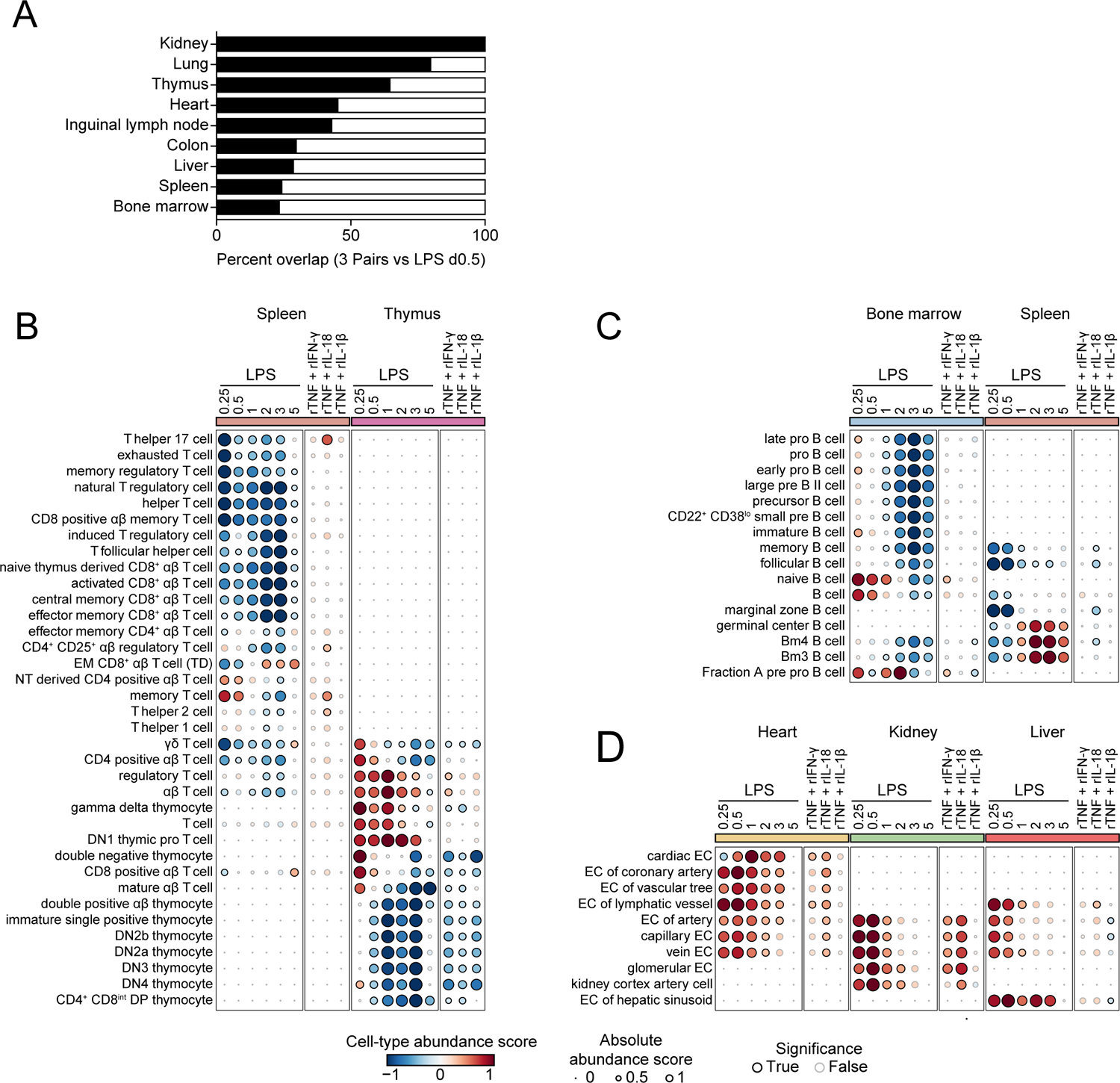
The Pairwise Effects of TNF plus IL-18, IFN-γ, or IL-1β Lead to Well-Known Sepsis Effects on Cells from Lymphoid and Non-Lymphoid Tissues, Related to Figure 4. (A) Percentages (black bars; x axis) of the effects of LPS on cell type abundance scores across tissues (y axis) mirrored by at least one of the three cytokine pairs tested: TNF plus IL-18, IFN-γ, or IL-1β. (B-D) Cell type abundance scores computed for indicated cell types (rows) and tissues (colors; top) upon injection of a sublethal dose of LPS in wild-type (left) or indicated knockout animals pre-treated with indicated neutralizing antibodies (right) or injected with indicated recombinant cytokine pairs (center) (columns). Black borders indicate significance (z-score > 1). EM, effector memory; TD, terminally differentiated; NT, naive thymus; DP, double positive; EC, epithelial cell.

**Figure S5.**
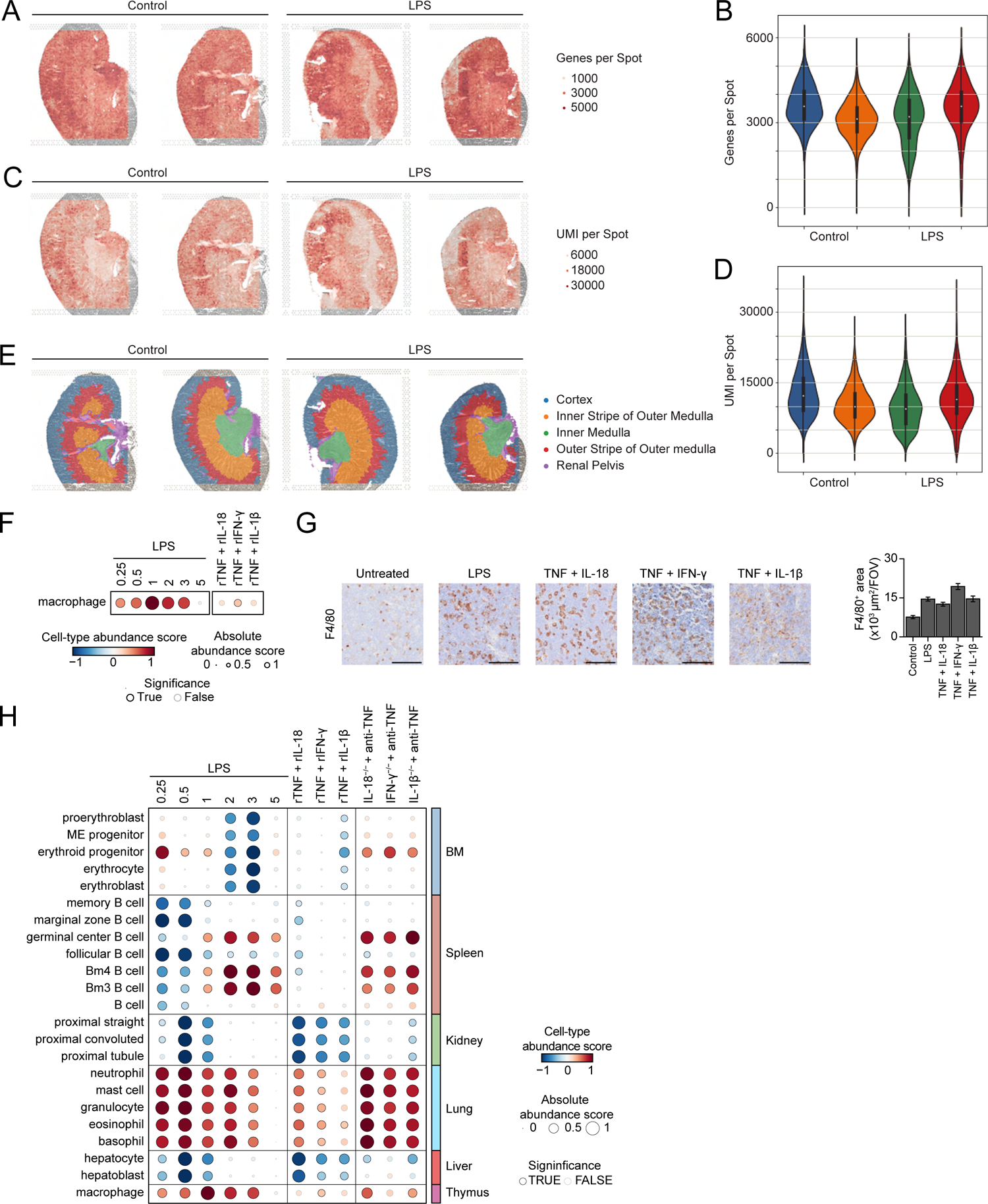
Experimental Validation of Changes in Cell Type Abundance Scores Computed from Whole-Tissue Gene Expression Profiles in Kidney and Thymus, Related to Figure 5. (A-E) Grey-scale H&E images from mouse kidney sections (n = 2) from PBS-(control) or LPS-treated mice processed for spatial transcriptomics and overlaid with the numbers of genes (A) or UMIs (C) detected per spot, or with spatial clusters annotated with known kidney histological regions (E). Violin plots show the matching distributions of the numbers of genes (B) and UMIs (D) per spot. (F) Cell type abundance scores computed for thymic macrophages (row) upon injection of a sublethal dose of LPS in wild-type (left) or injected with indicated recombinant cytokine pairs (right) (columns). (G) Images (40X magnification; left) from F4/80 immunohistochemistry in thymus tissues from mice injected with LPS, indicated cytokines, or left untreated as controls. Bar graph (right) shows quantifications of F4/80^+^ areas per field of view (FOV). Scale bars, 100µm; Error bar, SEM (n = 15). (H) Cell type abundance scores computed for thymic macrophages (row) upon injection of a sublethal dose of LPS in wild-type (left) or indicated knockout animals pre-treated with indicated neutralizing antibodies (right) or injected with indicated recombinant cytokine pairs (center) (columns).

**Figure S6.**
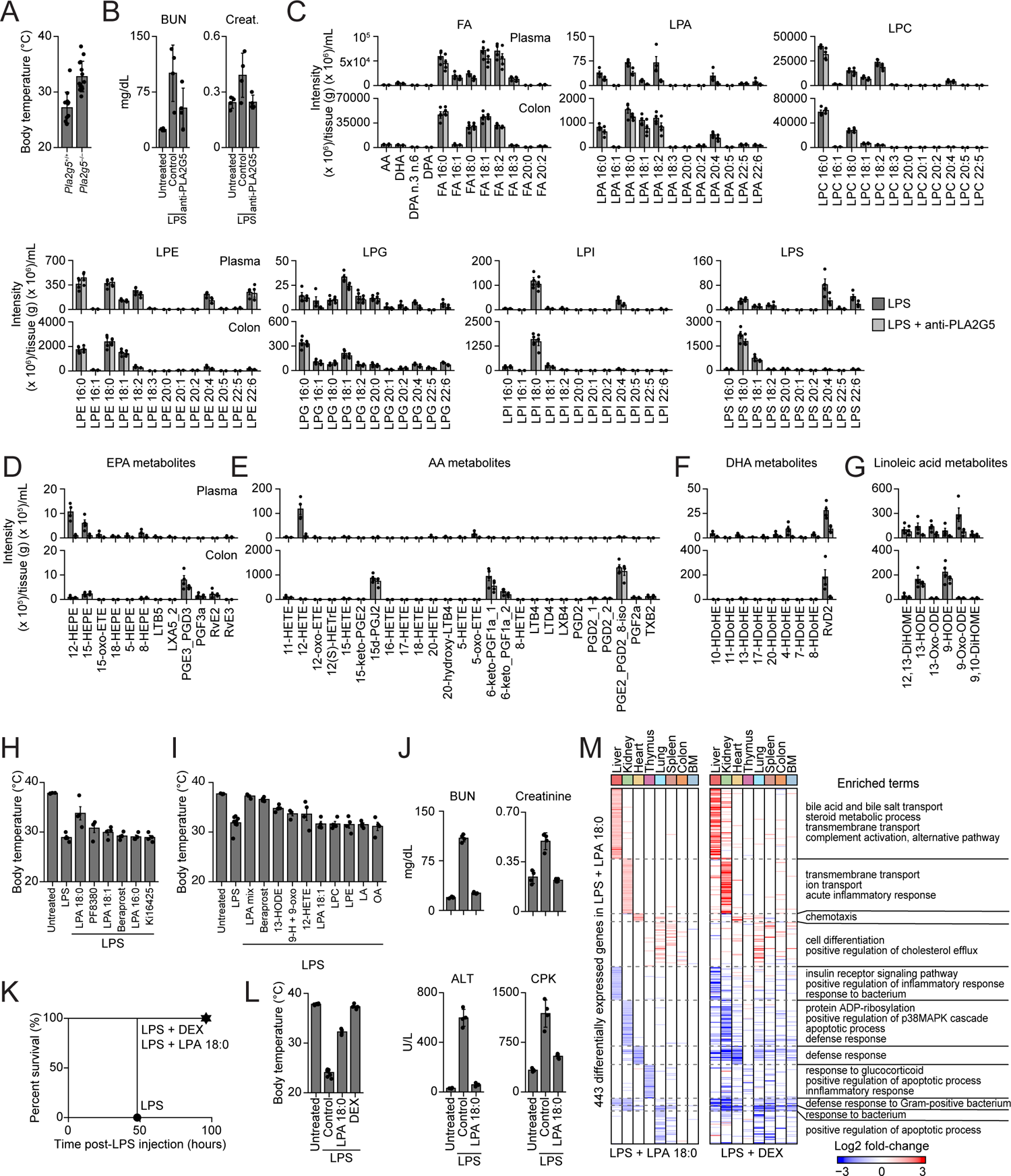
PLA2G5 Blockade Impacts the Plasma Levels of Several Lipid Metabolites, such as LPA 18:0 which Rescued Mice from Lethal Sepsis through Anti-Inflammatory Activities, Related to Figure 6. (A) Rectal temperature in *Pla2g5*^+/+^ or *Pla2g5*^-/-^ mice 24 hours after LPS injection. Error bars, SD (n = 12). (B) Serum levels of indicated organ injury markers before (control) and 12 hours post-injection with a sublethal LPS dose in the presence or absence of Pla2g5 blockade. Error bars, SD (n = 4). (C-G) Lipidomics profiling in plasma and colon tissues from mice after 12 hours post-sublethal LPS injection and pre-treated with anti-Pla2g5 neutralizing antibodies or left untreated. Shown are mass spectrometry intensities for indicated lipid metabolites: FA, fatty acid; LPA, lysophosphatidic acid; LPC, lysophosphatidylcholine; LPE, lysophosphatidylethanolamine; LPG, lysophosphatidylglycerol; LPI, lysophosphatidylinositol; LPS, lysophosphatidylserine; EPA, eicosapentaennoic acid; AA, arachidonic acid; DHA, docosahexaenoic acid; LA, linoleic acid, metabolites; PGE2_PGD2_8-iso, PGE2_PGD2_8-iso-PGA2; Error bar, SEM (n = 4). (H-I) Measurements of rectal temperature in mice untreated (control) or injected with a lethal (H) or sublethal (I) dose of LPS, at 9 and 12 hours post-LPS, respectively, and with pre-treatment with indicated lipid species or inhibitors (PF8380 or Ki16425), or without pre-treatment (LPS only control). LPA mix, LPA 16:0, 18:0, and 18:1; 9-H + 9-oxo, 9-HODE + 9-oxo-ODE. (J) Serum levels of indicated organ injury markers before (control) and 12 hours post-injection with a sublethal LPS dose in the presence or absence of pre-treatment with LPA 18:0. Error bars, SD (n = 4). (K-L) Measurements of survival (K) and rectal temperature (L; 24 hours post-LPS) in mice injected with a lethal dose of LPS in the presence or absence of pre-treatment with LPA 18:0 or dexamethasone (DEX). Error bars, SD (n = 5). (M) Heatmaps of differentially expressed genes (rows) from whole-tissue mRNA profiles of indicated organs (color; top) ordered by k-means clustering at 16 hours post-sublethal LPS injection in the presence of LPA 18:0 (left) or 12 hours post-sublethal LPS injection in the presence of DEX (right) pre-treatment. Values are log2 fold-changes relative to matching organs from LPS-injected mice without pre-treatment (FDR-adjusted p-value < 0.05; absolute fold change > 2; n = 4). Representative enriched Gene Ontology (GO) terms within each cluster are indicated on the right (p-value < 0.05).

**Figure S7.**
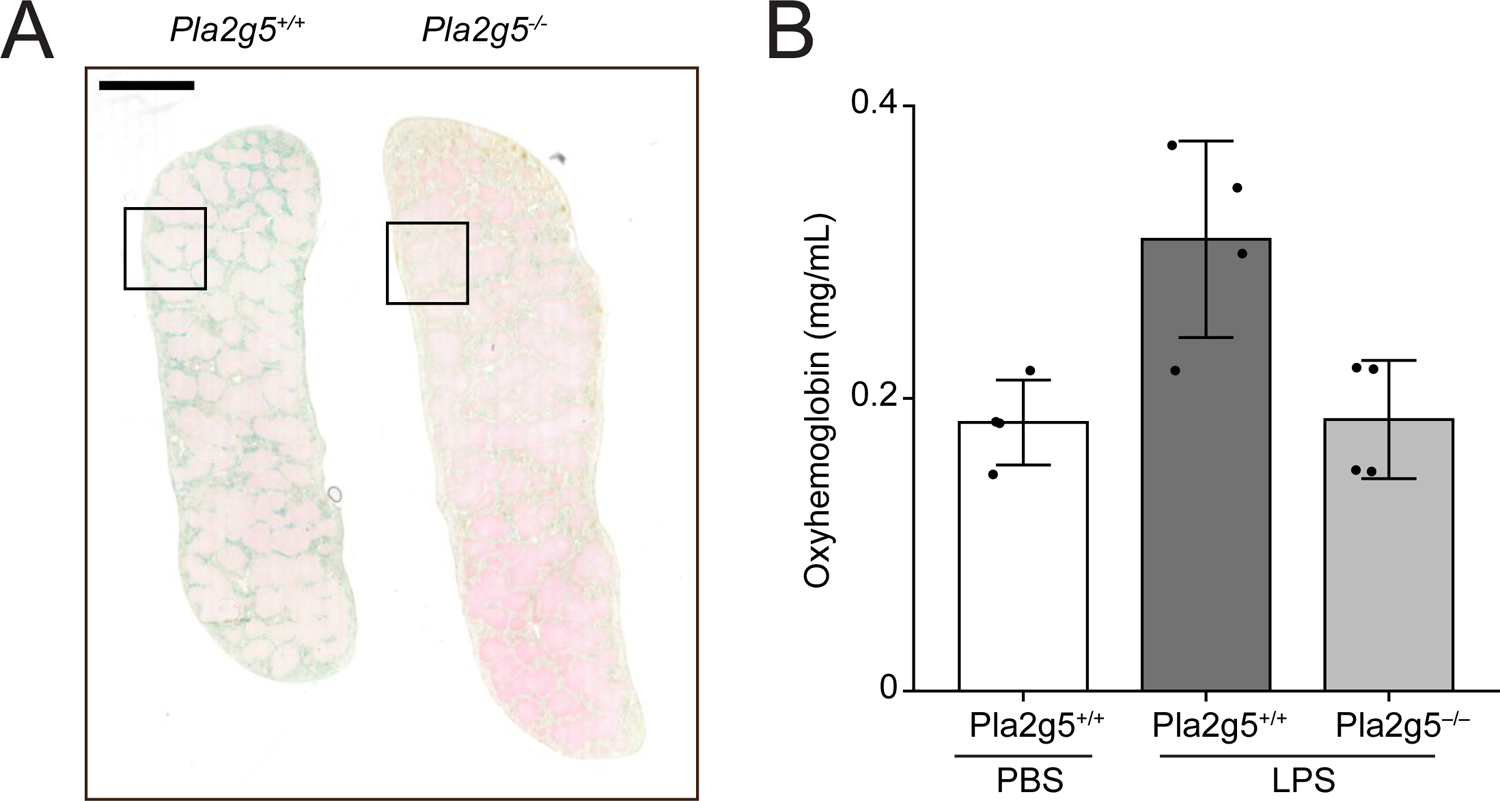
PLA2G5 Regulates Hemolysis in Blood and Impacts Iron Metabolism in Spleen, Related to Figure 7. (A) Images (40X magnification) of Prussian blue staining for ferric iron on spleen histological sections from LPS-injected *Pla2g5*^+/+^ (top) or *Pla2g5*^-/-^ (bottom) mice. Insets indicate the images shown in Figure 7F. Scale bar, 2 mm. (B) Plasma oxyhemoglobin levels in mice injected with plasma collected from LPS-injected *Pla2g5*^+/+^ or *Pla2g5*^-/-^ mice or PBS-injected *Pla2g5*^+/+^ mice as control. Error bars, SD (n = 4).

## SUPPLEMENTARY TABLES

**Table S1.** Whole-tissue RNA-seq analysis of LPS or CLP sepsis from wild-type, *Irf1^−/−^*, and *Stat1^−/−^* mice, related Figure 1. (A) Log2 fold-change in expression for all differentially expressed (DE) genes across the 13 organs measured from LPS-injected mice relative to controls (related to Figure 1C). (B) List of sepsis biomarker genes published by Pierrakos and colleagues (from reference 31). (C) Log2 fold-change in expression for all DE genes across indicated tissues from LPS-injected or CLP mice relative to controls (related to Figure S1D). (D) Enrichment analysis for genes in each topic (related to Figure 1F). (E) Log2 fold-change in expression for all DE genes across indicated tissues from LPS-injected wild-type, *Irf1^−/−^*, or *Stat1^−/−^* mice relative to control (related to Figure S1J).

**Table S2.** Whole-tissue RNA-seq analysis from mice injected with 6 singles or 15 pairs of recombinant cytokines, related to Figure 2. (A-B) Log2 fold-change in expression for all DE genes across indicated tissues from mice injected with 6 recombinant cytokines used alone (A) or in pairwise combinations (B) relative to untreated, control tissues (related to Figure S2A-B). (C) Log2 fold-change in expression for all DE genes across indicated tissues from LPS-injected mice (day 0.5) relative to controls (left), and row annotations indicating DE genes in at least one organ from mice injected with each of the 15 cytokine pairs tested (right) (related to Figure 2C).

**Table S3.** Synergistic and antagonistic transcriptional effects of 3 cytokine pairs, related to Figure 3. Log2 fold-change in expression for all DE genes across indicated tissues from mice injected with indicated recombinant cytokine pairs relative to controls (left), and annotations indicating genes showing significant synergistic or antagonistic differential expression in pairs relative to matching single cytokines (right) (related to Figure 3D).

**Table S4.** Impact of LPS and recombinant cytokine pairs on 195 cell types across 9 organ types, related to Figure 4. Relative cell abundance scores (Methods) across indicated tissues from wild-type mice injected with LPS at indicated time points or indicated recombinant cytokine pairs (related to Figure 4B).

**Table S5.** Lipidomics analysis of plasma and colon tissues from LPS-injected mice with or without PLA2G5 blockade, related to Figure S6. (A) Lipidomics data of indicated tissues from LPS-injected mice with or without pre-treatment with anti-PLA2G5 neutralizing antibodies (related to Figure S6C-G). (B) Log2 fold-change in expression for all DE genes across indicated tissues from LPS-treated mice which where pre-treated with LPA 18:0 (left) or dexamethasone (right) (related to Figure S6M).

**Table S6.** Effect of PLA2G5 blockade on whole-tissue RNA-seq in LPS-injected mice, related to Figure 7. Log2 fold change in expression for all DE genes across indicated tissues from LPS-injected mice with or without pre-treatment with anti-PLA2G5 neutralizing antibodies (related to Figure 7A).

## METHODS

### RESOURCE AVAILABILITY

#### Data and Code Availability

The sequencing (whole-tissue and spatial RNA-seq) and mass spectrometry (lipidomics) datasets generated during this study have been respectively deposited in the Gene Expression Omnibus and MassIVE repositories under accession numbers GSExx, GSEyy and GSEzz, and XX, respectively. All scripts and preprocessed datasets are publicly available at the following repository: https://github.com/chevrierlab/xx.

## EXPERIMENTAL MODEL AND SUBJECT DETAILS

### Mice

Female C57BL/6J mice (wild-type, stock 000664), B6.129S7-*Ifng^tm1Ts^*/J (Ifng KO, stock 002287), B6.129P2-*Il18^tm1Aki^*/J (Il18 KO, stock 004130), C57BL/6J-*Il1b^em2Lutzy^*/Mmjax (Il1b KO, stock 068082-JAX), B6.129S-*Tnf^tm1Gkl^*/J (Tnf KO, stock 005540), B6.129S2-*Irf1^tm1Mak^*/J (Irf1 KO, stock 002762), and B6.129S(Cg)-*Stat1^tmDlv^*/J (Stat1 KO, stock 012606) were obtained from the Jackson Laboratories. *Pla2g5^−/−^* mice were kindly provided by Steven Dudek (University of Illinois at Chicago, USA).^102, 103^ Animals were housed in specific pathogen-free and BSL2 conditions at The University of Chicago, and all experiments were performed in accordance with the US National Institutes of Health Guide for the Care and Use of Laboratory Animals and approved by The University of Chicago Institutional Animal Care and Use Committee.

## METHOD DETAILS

### Sepsis induction

For LPS endotoxemia, mice were injected intraperitoneally with either lethal (10-15 mg/kg) or sublethal (3-5 mg/kg) doses of LPS derived from Escherichia coli O55:B5 (Sigma-Aldrich) diluted in PBS. Dosing was established for each lot of LPS by *in vivo* titration. Cecal ligation and puncture (CLP) was performed as described by others^22^. Briefly, mice were anesthetized with isoflurane. A 1- to 2-cm midline laparotomy was performed, and the cecum was exposed. The cecum was ligated with 6-0 silk suture (Ethicon) and perforated as follows to vary disease severity: (1) mild sepsis: ligate at distal 33% position and perforate once with a 21-G needle; (2) moderate sepsis: ligate at distal 40% position and perforate twice with a 19-G needle; (3) severe sepsis: ligate immediately below the ileocecal valve and perforate twice with a 19-G needle. The cecum was tucked back into the peritoneum and gently squeezed to extrude a small amount of fecal content. The peritoneal wall was closed using absorbable suture. The skin was closed with surgical staples. To resuscitate animals, 1 mL of saline was injected subcutaneously. Mice were temporarily placed on a heating pad for recovery. Sham operated mice underwent the same procedure except that the cecum was neither tied nor perforated.

### Recombinant cytokine injections

C57BL/6J mice were injected intravenously with 2.5 µg of recombinant TNF, IL-1β, IL-6, IL-10, IL-18, or IFN-γ used alone (6 singles) or in pairwise combinations (15 pairs).

### Neutralizing antibody and drug treatments

For neutralizing antibodies, C57BL/6J or indicated knockout mice were injected intraperitoneally with 50 µg of anti-PLA2G5 (clone MCL-3G1, Cayman Chemical), anti-TNF (clone BE0058, BioXCell), anti-IL-18 (clone BE0237, BioXCell), anti-IFN-γ (clone BE0055, BioXCell), or anti-IL-1β (clone BE0246, BioXCell) neutralizing antibodies in 100 µl of PBS 1 hour prior to LPS injection. For dexamethasone treatment, mice were injected intraperitoneally with 7 mg/kg of dexamethasone diluted in 100 µl of PBS 1 hour prior to LPS injection.

### Plasma transfer

Whole blood was harvested from donor C57BL/6J or *Pla2g5^−/−^* mice injected with LPS 12 hours prior by cardiac puncture. Plasma was isolated using lithium heparin coated Vacutainer tubes (BD) and transferred intravenously (100 µL) to recipient C57BL/6J mice.

### Lipid metabolite treatments

Mice were injected intraperitoneally with the following lipid species, analogs, or inhibitors at indicated doses per animal in 100 µL of PBS + 1% fatty acid-free BSA: LPA 16:0, LPA 18:0 and LPA 18:1 (used alone or as a mixture; 100 µg each), 12-HETE (100 ng), 13-HODE (100 ng), a mixture of 9-HODE and 9-oxo-ODE (100 ng each), beraprost (1 µg), linoleic acid (6.25 mg), oleic acid (6.25 mg), lysophosphatidyl ethanolamine (LPE, 0.2 mg), lysophosphatidyl choline (LPC, 0.2 mg), autotaxin inhibitor PF8380 (200 µg), and LPA receptor antagonist Ki16425 (400 µg).

### Blood analysis

Mouse whole blood was harvested by cardiac puncture and plasma and serum were isolated using lithium heparin coated Microtainer blood collection tubes (BD 365965) and Microtainer blood collection tubes (BD 365978), respectively. For flow cytometric, bead-based immunoassays, plasma was diluted and processed using the LEGENDplex Mouse Inflammation Panel (BioLegend 740446) and Mouse Macrophage/Microglia Panel (BioLegend 740846) kits. Data were acquired on the NovoCyte flow cytometer (Acea Biosciences/Agilent) and analyzed using the LEGNEDplex software v8 (BioLegend). To measure tissue injury marker levels in sera, samples were processed using mass spectrometry (In Vivo Animal Core, University of Michigan) or with the following kits for BUN (BioAssay Systems DIUR-100), ALT (Cayman Chemical 700260), and troponin-I (Life Diagnostics CTNI-1-HS) levels according to the manufacturer’s instructions. Total plasma heme was measured with the 3,3’,5,5’ tetramethylbenzidine (TMB) peroxidase assay and oxyhemoglobin with a Nanodrop One (Thermoscientific) instrument.

### Tissue harvest

Tissues were harvested, frozen and stored as previously described^24, 25^. Mice were anesthetized with 2,2,2-tribromoethanol (250-500 mg/kg) and perfused transcardially with PBS containing 10 mM EDTA (to avoid signal contamination from blood in tissues). Prior to perfusion, blood was collected by cardiac puncture and stored on ice, and immediately after perfusion, tissues were placed in RNA-preserving solution (5.3 M ammonium sulfate, 25 mM sodium citrate, 20 mM EDTA) and kept at 4°C overnight prior to transfer at −80°C for storage. For each mouse, we harvested up to 13 tissues in total: lymph node (inguinal), flank skin, thymus, heart, lung, spleen, kidney, small intestine, colon, liver, brain, bone marrow (BM) and peripheral blood mononuclear cells (PBMCs). Small intestine and colon were cut longitudinally and washed extensively in PBS to completely remove feces contamination. Bone marrow cells were collected from femora and tibiae, stored overnight in RNA-preserving solution at 4°C, centrifuged at 5,000 g for 5 min at 4°C, and cell pellets were stored at −80°C.

### Whole-tissue RNA extraction

Whole-tissue RNA extraction was performed as described previously.^25^ Briefly, tissues stored in RNA-preserving solution were thawed and transferred to 2 mL tubes containing 700-1500 µL (depending on tissue) of PureZOL (Bio-Rad, 7326890) or homemade Trizol-like solution (38% phenol, 0.8 M guanidine thiocyanate, 0.4 M ammonium thiocyanate, 0.1 M sodium acetate, 5% glycerol). Tissues were lysed by adding 2.8-mm ceramic beads (OMNI International, 19-646) and running 1-3 cycles of 5-45 s at 3500 rpm on the PowerLyzer 24 (QIAGEN). For liver, brain, and small intestine samples, tissues were lysed with 3-5 mL using M tubes (Miltenyi biotec, 130-096-335) and running 1-4 cycles of the RNA_02.01 program on the gentleMACS Octo Dissociator (Miltenyi biotec). Next, lysates were processed in deep 96-well plates (USA Scientific 1896-2000) by adding chloroform for phase separation by centrifugation, followed by precipitation of total RNA in the aqueous phase using magnetic beads coated with silane (Dynabeads MyOne Silane; TermoFisher Scientific 37002D), buffer RLT (QIAGEN 79216), and ethanol. Genomic DNA contamination was removed by on-bead DNase I (ThermoFisher Scientific AM2239) treatment at 37 °C for 20 min. After washing steps with 80% ethanol, RNA was eluted from beads. This RNA extraction protocol was performed on the Bravo Automated Liquid Handling Platform (Agilent).^25^ Sample concentrations were measured using a Nanodrop One (Thermo Scientific). RNA quality was confirmed using a Tapestation 4200 (Agilent Technologies).

### RNA sequencing

For each tissue sample, full-length cDNA was synthesized in 20 µl final reaction volume containing the following: (1) 10 µl of 10 ng/µl RNA; (2) 1 µl containing 2 pmoles of a custom RT primer biotinylated in 5’ and containing sequences from 5’ to 3’ for the Illumina read 1 primer, a 6-bp sample barcode (up to 384), a 10-bp unique molecular identifier (UMI), and an anchored oligo(dT)_30_ for priming;^104^ and (3) 9 µl of RT mix containing 4 µl of 5X RT buffer, 1 µl of 10 mM dNTPs, 2 pmoles of template switching oligo (TSO), and 0.25 µl of Maxima H Minus Reverse Trascriptase (Thermo Scientific, EP0753). First, barcoded RT primers were added to RNA, which were then denatured at 72°C for 1 min and snap cooled on ice. Second, the RT mix was added, and plates were incubated at 42°C for 120 min. For each library, double stranded cDNA from up to 384 samples were pooled using DNA Clean & Concentrator-5 columns (Zymo Research, D4013), and residual RT primers were removed using exonuclease I (New England Biolabs, M0293). Full-length cDNAs were amplified with 5 to 8 cycles of single-primer PCR using the Advantage 2 PCR Kit (clontech 639206) and cleaned up using SPRIselect magnetic beads (Beckman Coulter B23318). cDNA was quantified with a Qubit dsDNA High Sensitivity Assay Kit (ThermoFisher Scientific 32851) and 50 ng of cDNA per pool of samples was tagmented using the Tagment DNA Enzyme I (Illumina 20034197) and amplified using the NEBNext Ultra II Q5 Master Mix (NEW ENGLAND BioLabs M0544L). Libraries were gel purified using 2% E-Gel EX Agarose Gels (ThermoFisher Scientific G402002), quantified with a Qubit dsDNA High Sensitivity Assay Kit (ThermoFisher Scientific Q32851) and a Tapestation 4200 (Agilent Technologies), and sequenced on the NextSeq 550 platform (Illumina).

### Spatial transcriptomics

Mouse kidneys were dissected from LPS-injected or control mice without transcardial perfusion and frozen in Optimal Cutting Temperature (OCT) media. 10 µm frozen tissue sections were cut with a CM1850 Cryostat (Leica) and mounted onto a Visium Spatial Gene Expression library preparation slide (10X Genomics). Samples were processed to generate spatial transcriptomics sequencing libraries according to the manufacturer’s instructions. In brief, sections were fixed in 100% methanol and stained with hematoxylin and eosin (H&E) reagents. H&E-stained sections were imaged using a CRi Panoramic MIDI Whole Slide Scanner with 20X magnification. Sections were then permeabilized with 0.1% pepsin in 0.01M HCl for 14 min at 37°C and processed for in-tissue reverse transcription followed by on-slide second strand synthesis. Resulting cDNA was used to construct sequencing libraries that were sequenced on the NextSeq 550 platform (Illumina), with 28 bases for Read 1 and 56 for Read 2 and at a depth of 78-114 million total reads per sample.

### Histology

Tissue processing, embedding, sectioning, immunohistochemistry using purified anti-mouse Ly6G (clone 1A8, BioLegend) and F4/80 (clone BM8, BioLegend) antibodies, and H&E, TUNEL (Terminal deoxynucleotidyl transferase dUTP nick end labeling), or PAS (Periodic Acid-Schiff) staining was performed by the Human Tissue Resource Center at the University of Chicago. To stain ferric iron in tissue sections, spleens were frozen in OCT using dry ice, sectioned (10 µm) using a cryostat (Leica CM1850), and stained with Perl’s Prussian blue (Sigma-Aldrich, P3289) and neutral red (Sigma-Aldrich, 72210). Section images were obtained using the Slideview VS200 Research Slide Scanner (Olympus). Image analysis and quantification (Ly6G^+^, TUNEL^+^, and F4/80^+^ areas) was performed using the ImageJ software (https://imagej.nih.gov/ij/).

### Flow cytometry

To analyze splenic B cells, total splenocytes were obtained by mashing spleens on 70-µm filters followed by red blood cell lysis (Lonza). To analysis red blood cell content in the bone marrow, total bone marrow cells were flushed out of femora and tibiae using PBS. Single-cell suspensions were stained in the presence of Fc receptor-blocking antibodies (anti-mouse CD16/32, clone 93) using the following antibodies (BioLegend): CD19-FITC (clone 1D3/CD19, 152403), B220-PerCP (clone RA3-6B2, 103233), CD93-PE (clone AA4.1, 136503), CD23-APC (clone B3B4, 101619), CD21-Pacific Blue (clone 7E9, 123413), Ter119-FITC (clone TER-119, 116205), CD45-APC-Cy7 (clone 30-F11, 103115). Cell viability was measured using Zombie Yellow Fixable Viability kit (423103) or DAPI. Flow cytometry data were acquired on the NovoCyte flow cytometer (Acea Biosciences/Agilent Technologies) and analyzed using the FlowJo software (BD).

### In vitro erythrocyte lysis assay

Mouse whole blood was spun at 1,000 g, the plasma and buffy coat were removed, and red blood cells were washed three times in Hepes-buffered saline. Subsequently, a portion of erythrocytes was incubated with 100 µM CaCl_2_ in Hepes-buffered saline for 3 min at 37 °C. Calcium ionophore (Cayman Chemical, A23187) was added to a final concentration of 4 µM, and the cells were further incubated for 10 min at 37 °C to induce membrane phospholipid (PS) scrambling. After that, 2 mM EDTA was added to stop the reaction. PS-exposing or untreated erythrocytes (10^9^ cells/mL) were incubated with 100 ng/mL of human recombinant PLA2G5 (Cayman chemical, 10009563) for 1 h at 37 °C in Hepes-buffered saline with 2 mM of calcium for the activity of PLA2G5. Hemolysis was measured by determining the amount of oxyhemoglobin in the supernatant of erythrocyte suspensions after centrifugation at 4,000 g. The absorbance at 414 nm in the supernatant of erythrocyte suspensions was measured using Nanodrop (Thermoscientific) and compared to that of a hemolyzed aliquot of the same erythrocyte suspension treated with Triton X-100.

### Lipidomics analysis

The procedures for lipidomics analysis using high-performance liquid chromatography coupled with electrospray tandem mass spectrometry (LC-ESI-MS/MS) were described previously.^105^ Briefly, C57BL/6J were injected intraperitoneally with 50 µg of anti-PLA2G5 neutralizing antibodies in 100 µl of PBS 1 hour prior to LPS injection. Twelve hours after LPS injection, mice were anesthetized with 2,2,2-tribromoethanol (250-500 mg/kg) and perfused transcardially with PBS containing 10 mM EDTA. Plasma was collected from blood prior to perfusion and frozen at −80°C. Immediately after perfusion, colon tissues were extensively washed in PBS and frozen by liquid nitrogen and kept at −80°C until the following procedures. Tissues were mechanically homogenized with the Precellys 24 homogenizer (Bertin Technologies, Montigny-le-Bretonneux, France) in methanol containing internal standards (500 pmol/sample of d4-labeled EPA, d5-labeled PGE2, LPC with a 17:0 fatty acyl chain (LPC17:0), and PC with two 14:0 fatty acyl chains (PE14:0-14:0)) and then incubated overnight at −20°C. For extraction of phospholipids and lysophospholipids, one-tenth of tissue lysates were added to 10 volumes of 20 mM Tris-HCl (pH 7.4) and were extracted using the method of Bligh and Dyer.^106^ For extraction of oxygenated fatty acid metabolites, nine-tenths of the tissue lysates were added to water (final methanol concentration of 10% (v/v)), and the lipids were extracted using an Oasis HLB cartridge (Waters, Milford, MA, USA). The samples were applied to a Kinetex C18 column (Kinetex C18, 2.1 x 150 mm, 1.7 µm particle; Phenomenex, Inc., Torrance, CA, USA) connected with ESI-MS/MS on a liquid chromatography (NexeraX2 system; Shimadzu Co., Kyoto, Japan) coupled with a 4000Q-TRAP quadrupole-linear ion trap hybrid mass spectrometer (AB Sciex, Framingham, MA, USA). For analyses of free fatty acids (FFAs), lysophospholipids (LPLs) and phospholipids, the samples were applied to the column and separated by a step gradient with mobile phase A (acetonitrile/methanol/water =1/1/1 (v/v/v) containing 5 mM phosphoric acid and 1 mM ammonium formate) and mobile phase B (2-propanol containing 5 µM phosphoric acid and 1 mM ammonium formate) at a flow rate of 0.2 mL/min at 50°C. For analyses of oxygenated fatty acid metabolites, the samples were applied to the column and separated using a step gradient including mobile phase C (water containing 0.1 % acetic acid) and mobile phase D (acetonitrile/methanol = 4/1 (v/v)) at a flow rate of 0.2 mL/ min at 45°C. Identification of phospholipids, LPLs, FFAs, and oxygenated PUFAs (polyunsaturated fatty acids) metabolites was conducted by multiple reaction monitoring (MRM) transition, and quantification was performed based on the peak area of the MRM transition and the calibration curve obtained with an authentic standard for each compound.^107^

## QUANTIFICATION AND STATISTICAL ANALYSIS

### RNA sequencing data analysis

Sequencing read files were processed to generate UMI (unique molecular identifier)^108^ count matrices using the python toolkit from the bcbio-nextgen project version 1.1.5 (https://bcbio-nextgen.readthedocs.io/en/latest/). In brief, reads were aligned to the mouse mm10 transcriptome with RapMap.^109^ Quality control metrics were compiled with a combination of FastQC (http:bioinformatics.babraham.ac.uk/projects/fastqc/), Qualimap, MultiQC (https://github.com/ewels/MultiQC).^110^,^111^ Samples were demultiplexed using barcodes stored in Read 1 (first 6 bases) and raw UMI count matrices were computed using UMIs stored in Read 1 (bases 7 to 16) (https://github.com/vals/umis).

Differential expression (DE) analysis was done using custom scripts in R (http://www.R-project.org). Raw count matrices were filtered to keep genes with at least 20 counts per million (cpm) or 5 UMIs in 2 samples and normalized across samples using the calcNormFactor function in edgeR.^112^ We identified genes with at least 2-fold expression difference and indicated Benjamini and Hochberg FDR adjusted p value by comparing treated tissues and matching control tissues using limma.

To assess the expression profiles of known sepsis biomarkers, we used a set of 258 genes reported as sepsis biomarkers by others.^31^ We defined the absolute average Log_2_ fold change of these 258 genes within each RNA-seq profile as the sepsis biomarker score.

Pathway enrichment analysis was done on differentially expressed genes from indicated hierarchical or k-means clusters using DAVID.^113, 114^ The differentially expressed genes in topic 9 were subjected to enrichment analysis with Enrichr (http://amp.pharm.mssm.edu/enrichr/), using TRRUST_Transcription_Factors_2019.

Heatmaps for RNA-seq data display the indicated numbers of transcripts and color intensities are determined by log_2_ fold change value for each heatmap. The rows of each heatmap were ordered by k-means clustering of log_2_ fold change values in R or Morpheus (https://software.broadinstitute.org/morpheus). All heatmaps were generated using ComplexHeatmap (https://github.com/jokergoo/ComplexHeatmap) and circlize (https://github.com/jokergoo/circlize) packages in R.^115, 116^

### Topic modeling of whole-tissue RNA-seq data

#### Fitting the topic model

We used fastTopics to fit a topic model to the UMI counts,^33, 117^ with K = 16 topics. fastTopics implements the following two-step approach to fit the topic model: (1) fit a non-negative matrix factorization based on a Poisson model (“Poisson NMF”);^118^ (2) recover maximum-likelihood estimates (MLEs) of the topic model parameters by a simple reparameterization.^117^

In detail, we took the following steps. First, we removed genes with very low expression (total UMI count ≤ 20). Therefore, the K = 16 topic model was fit to UMI counts for 364 samples and 28,209 genes. Second, we ran 20 expectation maximization (EM) updates, without extrapolation, to get close to an MLE solution (“prefitting phase”). This prefitting phase was implemented in R by calling fit_poisson_nmf from fastTopics with the following settings: numiter = 20, method = “em”, init.method = “random”, control = list(nc = 8). Third, we performed an additional 180 coordinate descent (CD) updates, with extrapolation, to improve the fit (“refinement phase”). This refinement phase was implemented by calling fit_poisson_nmf with the following settings: method = “scd”, numiter = 180, control = list(numiter = 4, nc = 8, extrapolate = TRUE), in which the model fit was initialized using the fit obtained from the prefitting phase. Finally, the topic model was recovered from the Poisson NMF model by calling function poisson2multinom. The convergence diagnostics suggested that, after a total of 200 iterations of the Poisson NMF optimization, the parameter estimates were close to an MLE; the change in log-likelihood between successive iterations was less than 1 × 10-5, and the largest residual in the first order (“Karush-Kuhn-Tucker”) conditions was less than 1.

Reassuringly, the estimated topics captured the predominant expression patterns, most of which identify the 13 tissues (the one exception was lymph node and spleen, which shared the same expression pattern). Two other topics (topics 1 and 6) captured variation specific to two tissues (liver and PBMCs), and one topic (topic 9) captured changes in expression over time that were shared across most tissues.

### Visualizing the topic proportions

The *n* × *K* matrix of topic proportions, *L*, where *n* denotes the number of RNA-seq samples and *K* is the number of topics, can be viewed as an embedding of the samples in a (*K* − 1) dimensional space. A simple way to visualize this embedding in 2-d (or 3-d) is to apply a nonlinear dimensionality reduction technique such as *t*-SNE^119^ to *L*. An alternative powerful approach, first suggested by Dey *et al*,^33^ is to visualize all *K* − 1 dimensions simultaneously using a Structure plot, which has been used with great success in population genetics.^120^ The Structure plot is essentially a stacked bar chart, in which bars correspond to samples (rows of *L*) and bar heights (in different colors, one for each topic) are determined by the topic proportions. To arrange the samples in the Structure plot, we first grouped the RNA-seq samples by tissue, then we ordered them within each tissue by time point.

### Differential expression analysis allowing for grades of membership

To annotate the topics, we used the grade-of-membership differential expression (GoM DE) analysis methods recently developed by the Stephens lab and implemented in the de_analysis function in the fastTopics package. In brief, the GoM DE analysis is conceptually similar to a standard DE analysis,^121, 122^ but extends the idea of comparing expression between groups by allowing the cells to have partial membership to multiple groups (here, the groups are the topics in the topic model). We called the de_analysis function with the following settings: shrink.method = “ash”, pseudocount = 0.1 and control = list(ns = 1e5, nc = 8). We performed a second DE analysis, with the same settings, after merging topics 1 and 5 (capturing variation in PBMC expression), 2 and 13 (bone marrow) and topics 6 and 14 (liver). The GoM DE analysis produces, for each gene *j* and topic *k*, estimates of differences in expression, and statistics quantifying support for these differences. In the de_analysis interface, expression differences are defined by the “least extreme” log-fold change (“l.e. LFC”), which is defined for gene *j* and topic *k* as the log-fold change that is the smallest in magnitude among topic pairs (*k*, *l*). After computing initial estimates, the GoM DE analysis (with shrink.method = “ash”) performs an adaptive shrinkage step,^123^ separately for each topic, to stabilize the l.e. LFC estimates. We used the posterior mean estimates, posterior standard errors, posterior z-scores (posterior mean/s.e.) and local false sign rates (*lfsr*) produced by the adaptive shrinkage step to report results of the GoM DE analysis. Note that the *lfsr* can be interpreted similarly to the *q*-value [qvalue2003, qvalue2003b], for example, although the *lfsr* tends to be more conservative than quantities such as the *q*-value that control for the false discovery rate.^123^

### Gene set enrichment analysis

Mouse gene sets for the gene set enrichment analyses (GSEA) were compiled from the following gene set databases: NCBI BioSystems;^124^ Pathway Commons;^125, 126^ and MSigDB,^127–129^ which included Gene Ontology (GO) gene sets.^130, 131^ Specifically, we downloaded bsid2info.gz and biosystems_gene.gz from the NCBI FTP site (https://ftp.ncbi.nih.gov/gene) on March 22, 2020; PathwayCommons12.All.hgnc.gmt.gz from the Pathway Commons website (https://www.pathwaycommons.org) on March 20, 2020; and msigdb_v7.2.xml.gz from the MSigDB website (https://www.gsea-msigdb.org) on October 15, 2020. For the gene set enrichment analyses, we also downloaded the mouse gene information (“gene info”) file Mus_musculus.gene_info.gz from the NCBI FTP site on October 15, 2020. To facilitate integration of these gene sets into our analyses, we have compiled these gene sets into an R package (https://github.com/stephenslab/pathways). We performed two gene set enrichment analyses. In the first GSEA, we included all gene sets other than the following MSigDB collections: C1, C3, C4 and C6, and gene sets labeled as “archived”. In the second GSEA, we focused on the curated pathways, specifically gene sets belonging to the GO and CP subcategories in the MSigDB C2 gene set collection. In both analyses, we removed gene sets with fewer than 10 genes and with more than 400 genes. After removing these gene sets, 21,442 candidate gene sets remained for the first analysis, and 8,939 gene sets remained for the second analysis.

We took a simple multiple linear regression approach to the gene set enrichment analysis (GSEA), in which we modeled, for a given topic k, the (least extreme) LFC estimates for all genes using a multiple linear regression model in which the regression variables were gene-set memberships, 1 if gene i belongs to gene set j, otherwise 0. The model fitting for the multiple linear regression model was implemented in SuSiE.^132^ The idea behind this simple multiple linear regression approach is that the most relevant gene sets are those that best explain the log-fold changes, and therefore in the multiple regression we sought to identify these gene sets by finding regression coefficients that are nonzero with high probability. Modeling the LFC estimates also helped distinguish among DE genes that show only a slight increase in expression versus those that are highly overexpressed. This simple multiple linear approach ignored uncertainty in the LFC estimates. So, to address this issue, we shrunk the LFC estimates prior to running the GSEA; that is, we defined the regression outcome to be the posterior mean LFC estimate after applying adaptive shrinkage, as described above. This had the effect that genes that we were more uncertain about had an LFC estimate that was zero or near zero. A benefit to using SuSiE is that it automatically organizes similar or redundant gene sets into credible sets (CSs),^132^ making it easier to quickly recognize complementary gene sets.

In detail, the GSEA was performed as follows. We performed a separate GSEA for each topic, Specifically, for topic k, we ran the susieR function susie with the following options: L = 10, intercept = TRUE, standardize = FALSE, estimate_residual_variance = TRUE, refine = FALSE}, compute_univariate_zscore = FALSE and min_abs_corr = 0. We set L = 10 so that SuSiE returned at most 10 credible sets—that is, at most 10 enriched gene sets. For a given topic k, we reported a gene set as being enriched if it was included in at least one CS. We organized the enriched gene sets by credible set (specifically, 95% credible sets). We also recorded the Bayes factor for each CS, which gives a measure of the level of support for that CS. For each gene set included in a CS, we reported the posterior inclusion probability (PIP), and the posterior mean estimate of the regression coefficient. In the results, we refer to the regression coefficient as the enrichment coefficient for a given gene set j since it is an estimate of the expected increase in the LFC for genes that belong to gene set j relative to genes that do not belong to the gene set. Often, a CS contained only one gene set, in which case the PIP for that gene set was close to 1. In several other cases, the CS contained multiple similar gene sets; in these cases, the smaller PIPs indicate that it is difficult to choose among the gene sets because they are similar to each other. (Note that the sum of the PIPs in a 95% CS should always be above 0.95 and less than 1.) Occasionally, SuSiE returned a CS with a small Bayes factor containing a large number of gene sets. When this happened, we did not include these gene sets in the results.

### Computing environment for topic modeling and gene set enrichment analyses

Most computations on real data sets were performed in R 3.5.1 (https://www.R-project.org), linked to the OpenBLAS 0.2.19 optimized numerical libraries, on Linux machines (Scientific Linux 7.4) with Intel Xeon E5-2680v4 (“Broadwell”) processors. For performing the model topic model optimization and DE analysis, which included multithreaded computations, as many as 8 CPUs and 16 GB of memory were used.

### Statistical modeling of cytokine pairwise effects on tissue gene expression

To assess the extent to which pairwise administration of cytokines (*i.e.*, TNF plus IL18, IFN-γ, or IL-1β) resulted in non-additive changes (*i.e.*, synergistic or antagonistic interactions) in gene expression levels across tissues in mice, we developed an interaction scoring method based on a linear modeling method adapted from previous work,^133^ using custom scripts in R (https://www.r-project.org/).

First, raw, tissue RNA-seq count matrices were normalized across samples using the calcNormFactor function in edgeR^112^ and subsequently filtered to keep genes with at least 15 counts per million (cpm) in 2 samples. Data was log-transformed, and a linear model was fit using the limma package.^134^ We then computed the following contrasts for each pair (AB) of interest and its component singles (A, B), and unstimulated control mice:

Single 1 Effect = A – control,

Single 2 Effect = B – control,

Pair Effect = AB – control,

Additive Effect = [A – control] + [B – control], and

Interaction Effect = (AB – control) – [(A– control) + (B – control)].

Where, “Pair Effect” is equivalent to the observed gene expression value for a given pair, while “Additive Effect” is equivalent to the predicted gene expression value for that pair if it is assumed to be equal to the sum of the component singles. The “Interaction Effect” is equal to the difference between these two values and is used as the score for assessing non-additive interactions.

We identified genes with significantly different expression within each contrast and across all contrasts using a Benjamini and Hochberg correction for multiple hypothesis testing and an FDR of 0.1. We next classified each gene, for each organ and pair treatment, as “synergistic”, “antagonistic” or “additive”, depending on its score, and the gene expression values of the pair and its component singles. Genes without > 0.5 absolute difference in log2-fold change (LFC) in at least two of the three experimentally measured treatment conditions for a given pair and organ (Single 1 Effect, Single 2 Effect, Pair Effect) were considered to have roughly the same expression across all samples and were excluded from further classification to avoid classifying genes with very high or low baseline expression in the singles, (and therefore very high or low predicted additive effects but no additional increase or decrease in gene expression at the pair level) as synergistic or antagonistic.

Using the standard deviation for each contrast determined via limma, we calculated an error value, E, for each gene, as the average of the standard deviations for all experimentally measured samples for that gene. Where the score was > 2 * E, AND the score was significant (FDR < 0.1) OR score > 1 (> 2-fold difference between predicted and observed gene expression values), the gene was classified as synergistic. The gene was only classified as significantly synergistic if the score was determined to be significant at the chosen FDR (0.1). Following the same logic, if score < −2 * E & score significant (FDR < 0.1) OR score < −1, the gene was classified as antagonistic in a particular pair and organ. Again, only if the score was determined to be significant at the chosen FDR (0.1), was the gene classified as significantly antagonistic.

The total number of differentially expressed genes (DEGs) was calculated by totaling any gene which showed significant differential expression (FDR < 0.1) in single 1, single 2, or pair, compared to control. The percent of all differentially expressed genes (DEGs) for a given pair and organ that were classified as either synergistic, additive, or antagonistic was then calculated.

### Computing cell type abundance scores from whole-tissue RNA-seq profiles

#### Database preparation

We used the CellKb database (https://www.cellkb.com),135 which consists of marker gene sets for mouse cell types. The marker gene sets in CellKb are manually extracted from supplementary materials and raw or processed data from publications describing single-cell or bulk RNA-seq experiments and other publicly available databases, including Tabula Muris (https://tabula-muris.ds.czbiohub.org/), MSigDB (https://www.gsea-msigdb.org/gsea/msigdb/), and the Human Protein Atlas (proteinatlas.org).^42, 128, 129, 136, 137^ Marker gene sets are extensively curated to identify valid genes and cell types. Each marker gene set is associated with the cell type, tissue, organ and/or disease condition given in the source publication, which are mapped to standardized ontology terms using Cell Ontology, Uberon Ontology, and Disease Ontology.^138–140^ The quality of the marker gene sets for immune cell types has been evaluated by a comparison with the ImmGen database (https://www.immgen.org/).135

#### Calculation of cell type specificity scores for each gene

A cell type specificity score is calculated for each gene in CellKb based on its specificity and prevalence in marker gene sets of mouse cell types. It defines how specific a gene is for a given cell type and varies between 0 (lowest specificity) and 1 (highest specificity). For this study, we calculated cell type specificity scores for each gene across all mouse cell types using 10,614 marker gene sets extracted from 373 publications as follows:

Cell type specificity of gene i in cell type j, c_ij_ = max(0, r_wt_)

Where,

- Weighted ranked of gene i, 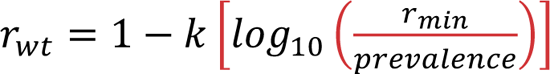
- r_min_ = min(rank of gene i in all marker gene sets of cell type j)
- prevalence = number of marker gene sets of cell type j where rank of gene i is ≤ 500
- constant k = 0.3
- normalized cell type specificity of gene i for cell type j, 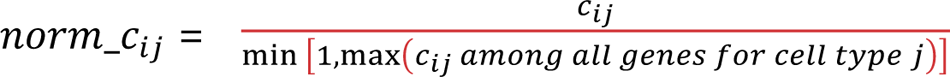

#### Calculation of relative cell type abundance scores for each cell type in each sample

To identify the cell types whose abundance is changing across tissues and disease conditions, a cell type activity score was calculated for each cell type in each whole-tissue RNA-seq sample as follows:

Cell type activity score for cell type j in condition k, 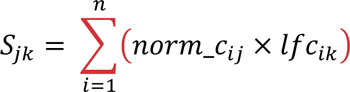

Where,

- norm_c_ij_ = normalized cell type specificity of gene i for cell type j
- lfc_ik_ = log fold change of gene i in condition k
- normalized cell type activity score for cell type j in condition k, 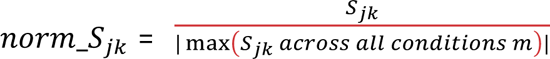

Z-scores were calculated for each cell type in each condition across all cell type abundance scores as follows and cell types above an absolute z-score of 1 were considered significant.

Zscore for cell type activity score of cell type j in condition k, 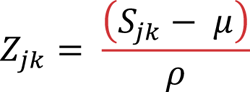

Where,

- S_jk_ = cell type activity score of cell type j in condition k
- μ = mean cell type activity score of all cell types across all conditions
- ρ = standard deviation of cell type activity score of all cell types across all conditions

### Spatial Transcriptomics

The output data of each sequencing run (Illumina BCL files) were converted into FASTQ files using Bcl2Fastq v2.19.1. The Space Ranger software (v1.2.0, 10X Genomics) was used to process, align, and summarize the FASTQ files against a GRCm39 mouse reference genome. Raw UMI (unique molecular identifier) count spot matrices, spot coordinates, and images were imported into python using Scanpy v1.9.1.^141^ Raw UMI counts were log10 normalized and clustered using a Louvain algorithm (resolution = 0.35). Differential expression between control and LPS-treated samples was performed using Scanpy’s rank_genes_groups function using a Wilcoxon rank-sum test. Spatially resolved counts of differentially expressed genes were overlayed with corresponding grey-scale H&E images and visualized using seaborn v0.11.2 (https://github.com/mwaskom/seaborn).

### Public RNA-sequencing data

To assess the expression pattern of *Pla2g5* across the body, we obtained single-cell RNA-seq data from the Tabula Muris Senis website (https://figshare.com/projects/Tabula_Muris_Senis/64982) and used the package TabulaMurisSenisData (github.com/fmicompbio/TabulaMurisSenisData) for data visualization. To compare the expression profile of bacterial sepsis (this study) with that of viral sepsis induced using tissues from mice infected with a lethal dose of Vaccinia virus strain Western Reserve,^24^ we used our previously published bulk RNA-seq data (GSE87633).

## REFERENCES

1. Marshall, J.C. (2018). Sepsis Definitions: A Work in Progress. Critical Care Clinics 34, 1–14. 10.1016/j.ccc.2017.08.004.

2. Singer, M., Deutschman, C.S., Seymour, C.W., Shankar-Hari, M., Annane, D., Bauer, M., Bellomo, R., Bernard, G.R., Chiche, J.-D., Coopersmith, C.M., et al. (2016). The Third International Consensus Definitions for Sepsis and Septic Shock (Sepsis-3). JAMA 315, 801. 10.1001/jama.2016.0287.

3. Marshall, J.C. (2014). Why have clinical trials in sepsis failed? Trends in Molecular Medicine 20, 195–203. 10.1016/j.molmed.2014.01.007.

4. Reinhart, K., Daniels, R., Kissoon, N., Machado, F.R., Schachter, R.D., and Finfer, S. (2017). Recognizing Sepsis as a Global Health Priority — A WHO Resolution. N Engl J Med 377, 414–417. 10.1056/NEJMp1707170.

5. Cavaillon, J.-M., Singer, M., and Skirecki, T. (2020). Sepsis therapies: learning from 30 years of failure of translational research to propose new leads. EMBO Molecular Medicine 12, e10128. 10.15252/emmm.201810128.

6. van der Poll, T., van de Veerdonk, F.L., Scicluna, B.P., and Netea, M.G. (2017). The immunopathology of sepsis and potential therapeutic targets. Nat Rev Immunol 17, 407–420. 10.1038/nri.2017.36.

7. Rudd, K.E., Johnson, S.C., Agesa, K.M., Shackelford, K.A., Tsoi, D., Kievlan, D.R., Colombara, D.V., Ikuta, K.S., Kissoon, N., Finfer, S., et al. (2020). Global, regional, and national sepsis incidence and mortality, 1990-2017: analysis for the Global Burden of Disease Study. Lancet 395, 200–211. 10.1016/S0140-6736(19)32989-7.

8. Rubio, I., Osuchowski, M.F., Shankar-Hari, M., Skirecki, T., Winkler, M.S., Lachmann, G., La Rosée, P., Monneret, G., Venet, F., Bauer, M., et al. (2019). Current gaps in sepsis immunology: new opportunities for translational research. The Lancet Infectious Diseases 19, e422–e436. 10.1016/S1473-3099(19)30567-5.

9. van der Poll, T., Shankar-Hari, M., and Wiersinga, W.J. (2021). The immunology of sepsis. Immunity 54, 2450–2464. 10.1016/j.immuni.2021.10.012.

10. Wiersinga, W.J., Leopold, S.J., Cranendonk, D.R., and van der Poll, T. (2014). Host innate immune responses to sepsis. Virulence 5, 36–44. 10.4161/viru.25436.

11. Hotchkiss, R.S., Moldawer, L.L., Opal, S.M., Reinhart, K., Turnbull, I.R., and Vincent, J.-L. (2016). Sepsis and septic shock. Nat Rev Dis Primers 2, 1–21. 10.1038/nrdp.2016.45.

12. Angus, D.C., and van der Poll, T. (2013). Severe sepsis and septic shock. N Engl J Med 369, 2063. 10.1056/NEJMc1312359.

13. Fajgenbaum, D.C., and June, C.H. (2020). Cytokine Storm. New England Journal of Medicine 383, 2255–2273. 10.1056/NEJMra2026131.

14. Callard, R., George, A.J.T., and Stark, J. (1999). Cytokines, Chaos, and Complexity. Immunity 11, 507–513. 10.1016/S1074-7613(00)80125-9.

15. Hotchkiss, R.S., and Nicholson, D.W. (2006). Apoptosis and caspases regulate death and inflammation in sepsis. Nat Rev Immunol 6, 813–822. 10.1038/nri1943.

16. Hotchkiss, R.S., Monneret, G., and Payen, D. (2013). Sepsis-induced immunosuppression: from cellular dysfunctions to immunotherapy. Nat Rev Immunol 13, 862–874. 10.1038/nri3552.

17. Hotchkiss, R.S., Swanson, P.E., Freeman, B.D., Tinsley, K.W., Cobb, J.P., Matuschak, G.M., Buchman, T.G., and Karl, I.E. (1999). Apoptotic cell death in patients with sepsis, shock, and multiple organ dysfunction. Critical Care Medicine 27, 1230.

18. Shankar-Hari, M., Fear, D., Lavender, P., Mare, T., Beale, R., Swanson, C., Singer, M., and Spencer, J. (2017). Activation-Associated Accelerated Apoptosis of Memory B Cells in Critically Ill Patients With Sepsis. Critical Care Medicine 45, 875. 10.1097/CCM.0000000000002380.

19. Brown, K., Brain, S., Pearson, J., Edgeworth, J., Lewis, S., and Treacher, D. (2006). Neutrophils in development of multiple organ failure in sepsis. The Lancet 368, 157–169. 10.1016/S0140-6736(06)69005-3.

20. Delano, M.J., and Ward, P.A. (2016). The immune system’s role in sepsis progression, resolution, and long-term outcome. Immunological Reviews 274, 330–353. 10.1111/imr.12499.

21. Lewis, A.J., Seymour, C.W., and Rosengart, M.R. (2016). Current Murine Models of Sepsis. Surg Infect (Larchmt) 17, 385–393. 10.1089/sur.2016.021.

22. Rittirsch, D., Huber-Lang, M.S., Flierl, M.A., and Ward, P.A. (2009). Immunodesign of experimental sepsis by cecal ligation and puncture. Nat Protoc 4, 31–36. 10.1038/nprot.2008.214.

23. Sjaastad, F.V., Jensen, I.J., Berton, R.R., Badovinac, V.P., and Griffith, T.S. (2020). Inducing Experimental Polymicrobial Sepsis by Cecal Ligation and Puncture. Current Protocols in Immunology 131, e110. 10.1002/cpim.110.

24. Kadoki, M., Patil, A., Thaiss, C.C., Brooks, D.J., Pandey, S., Deep, D., Alvarez, D., von Andrian, U.H., Wagers, A.J., Nakai, K., et al. (2017). Organism-Level Analysis of Vaccination Reveals Networks of Protection across Tissues. Cell 171, 398–413.e21. 10.1016/j.cell.2017.08.024.

25. Pandey, S., Takahama, M., Gruenbaum, A., Zewde, M., Cheronis, K., and Chevrier, N. (2020). A whole-tissue RNA-seq toolkit for organism-wide studies of gene expression with PME-seq. Nat Protoc, 1–25. 10.1038/s41596-019-0291-y.

26. McBride, M.A., Patil, T.K., Bohannon, J.K., Hernandez, A., Sherwood, E.R., and Patil, N.K. (2021). Immune Checkpoints: Novel Therapeutic Targets to Attenuate Sepsis-Induced Immunosuppression. Frontiers in Immunology 11.

27. Bateman, R.M., Sharpe, M.D., Singer, M., and Ellis, C.G. (2017). The Effect of Sepsis on the Erythrocyte. Int J Mol Sci 18, 1932. 10.3390/ijms18091932.

28. Kato, H., Itoh-Nakadai, A., Matsumoto, M., Ishii, Y., Watanabe-Matsui, M., Ikeda, M., Ebina-Shibuya, R., Sato, Y., Kobayashi, M., Nishizawa, H., et al. (2018). Infection perturbs Bach2- and Bach1-dependent erythroid lineage ‘choice’ to cause anemia. Nature Immunology 19, 1059–1070. 10.1038/s41590-018-0202-3.

29. Jensen, I.J., Sjaastad, F.V., Griffith, T.S., and Badovinac, V.P. (2018). Sepsis-Induced T Cell Immunoparalysis: The Ins and Outs of Impaired T Cell Immunity. The Journal of Immunology 200, 1543–1553. 10.4049/jimmunol.1701618.

30. Hoyer, F.F., Naxerova, K., Schloss, M.J., Hulsmans, M., Nair, A.V., Dutta, P., Calcagno, D.M., Herisson, F., Anzai, A., Sun, Y., et al. (2019). Tissue-Specific Macrophage Responses to Remote Injury Impact the Outcome of Subsequent Local Immune Challenge. Immunity 51, 899–914.e7. 10.1016/j.immuni.2019.10.010.

31. Pierrakos, C., Velissaris, D., Bisdorff, M., Marshall, J.C., and Vincent, J.-L. (2020). Biomarkers of sepsis: time for a reappraisal. Critical Care 24, 287. 10.1186/s13054-020-02993-5.

32. Lelubre, C., and Vincent, J.-L. (2018). Mechanisms and treatment of organ failure in sepsis. Nat Rev Nephrol 14, 417–427. 10.1038/s41581-018-0005-7.

33. Dey, K.K., Hsiao, C.J., and Stephens, M. (2017). Visualizing the structure of RNA-seq expression data using grade of membership models. PLOS Genetics 13, e1006599. 10.1371/journal.pgen.1006599.

34. Daix, T., Guerin, E., Tavernier, E., Mercier, E., Gissot, V., Hérault, O., Mira, J.-P., Dumas, F., Chapuis, N., Guitton, C., et al. (2018). Multicentric Standardized Flow Cytometry Routine Assessment of Patients With Sepsis to Predict Clinical Worsening. Chest 154, 617– 627. 10.1016/j.chest.2018.03.058.

35. Sadler, A.J., and Williams, B.R.G. (2008). Interferon-inducible antiviral effectors. Nat Rev Immunol 8, 559–568. 10.1038/nri2314.

36. Senaldi, G., Shaklee, C.L., Guo, J., Martin, L., Boone, T., Mak, T.W., and Ulich, T.R. (1999). Protection Against the Mortality Associated with Disease Models Mediated by TNF and IFN-γ in Mice Lacking IFN Regulatory Factor-1. The Journal of Immunology 163, 6820– 6826.

37. Karaghiosoff, M., Steinborn, R., Kovarik, P., Kriegshäuser, G., Baccarini, M., Donabauer, B., Reichart, U., Kolbe, T., Bogdan, C., Leanderson, T., et al. (2003). Central role for type I interferons and Tyk2 in lipopolysaccharide-induced endotoxin shock. Nat Immunol 4, 471–477. 10.1038/ni910.

38. Ince, C., Mayeux, P.R., Nguyen, T., Gomez, H., Kellum, J.A., Ospina-Tascón, G.A., Hernandez, G., Murray, P., De Backer, D., and Workgroup, on behalf of the A.X. (2016). The Endothelium in Sepsis. Shock 45, 259. 10.1097/SHK.0000000000000473.

39. Taylor, M.D., Brewer, M.R., Nedeljkovic-Kurepa, A., Yang, Y., Reddy, K.S., Abraham, M.N., Barnes, B.J., and Deutschman, C.S. (2020). CD4 T Follicular Helper Cells Prevent Depletion of Follicular B Cells in Response to Cecal Ligation and Puncture. Frontiers in Immunology 11.

40. Murakami, M., Sato, H., and Taketomi, Y. (2020). Updating Phospholipase A2 Biology. Biomolecules 10, 1457. 10.3390/biom10101457.

41. Samuchiwal, S.K., and Balestrieri, B. (2019). Harmful and protective roles of group V phospholipase A2: Current perspectives and future directions. Biochimica et Biophysica Acta (BBA) - Molecular and Cell Biology of Lipids 1864, 819–826. 10.1016/j.bbalip.2018.10.001.

42. Almanzar, N., Antony, J., Baghel, A.S., Bakerman, I., Bansal, I., Barres, B.A., Beachy, P.A., Berdnik, D., Bilen, B., Brownfield, D., et al. (2020). A single-cell transcriptomic atlas characterizes ageing tissues in the mouse. Nature, 1–6. 10.1038/s41586-020-2496-1.

43. Muñoz, N.M., Kim, K.P., Han, S.K., Boetticher, E., Sperling, A.I., Sano, H., Han, K., Zhu, X., Leff, A.R., and Cho, W. (2000). Characterization of Monoclonal Antibodies Specific for 14-kDa Human Group V Secretory Phospholipase A _2_ (hVPLA _2_). Hybridoma 19, 171– 176. 10.1089/02724570050031220.

44. Murch, O., Collin, M., and Thiemermann, C. (2007). LYSOPHOSPHATIDIC ACID REDUCES THE ORGAN INJURY CAUSED BY ENDOTOXEMIA-A ROLE FOR G-PROTEIN-COUPLED RECEPTORS AND PEROXISOME PROLIFERATOR-ACTIVATED RECEPTOR-γ. Shock 27, 48–54. 10.1097/01.shk.0000235086.63723.7e.

45. Horby, P., Lim, W.S., Emberson, J.R., Mafham, M., Bell, J.L., Linsell, L., Staplin, N., Brightling, C., Ustianowski, A., Elmahi, E., et al. (2021). Dexamethasone in Hospitalized Patients with Covid-19. N Engl J Med 384, 693–704. 10.1056/NEJMoa2021436.

46. Gibbison, B., López-López, J.A., Higgins, J.P.T., Miller, T., Angelini, G.D., Lightman, S.L., and Annane, D. (2017). Corticosteroids in septic shock: a systematic review and network meta-analysis. Critical Care 21, 78. 10.1186/s13054-017-1659-4.

47. Kohyama, M., Ise, W., Edelson, B.T., Wilker, P.R., Hildner, K., Mejia, C., Frazier, W.A., Murphy, T.L., and Murphy, K.M. (2009). Role for Spi-C in the development of red pulp macrophages and splenic iron homeostasis. Nature 457, 318–321. 10.1038/nature07472.

48. Haldar, M., Kohyama, M., So, A.Y.-L., Kc, W., Wu, X., Briseño, C.G., Satpathy, A.T., Kretzer, N.M., Arase, H., Rajasekaran, N.S., et al. (2014). Heme-Mediated SPI-C Induction Promotes Monocyte Differentiation into Iron-Recycling Macrophages. Cell 156, 1223–1234. 10.1016/j.cell.2014.01.069.

49. Garcia-Garcia, H.M., and Serruys, P.W. (2009). Phospholipase A2 inhibitors. Current Opinion in Lipidology 20, 327. 10.1097/MOL.0b013e32832dd4c7.

50. Larsen, R., Gozzelino, R., Jeney, V., Tokaji, L., Bozza, F.A., Japiassú, A.M., Bonaparte, D., Cavalcante, M.M., Chora, Â., Ferreira, A., et al. (2010). A Central Role for Free Heme in the Pathogenesis of Severe Sepsis. Science Translational Medicine 2, 51ra71–51ra71. 10.1126/scitranslmed.3001118.

51. Larsen, R., Gouveia, Z., Soares, M., and Gozzelino, R. (2012). Heme Cytotoxicity and the Pathogenesis of Immune-Mediated Inflammatory Diseases. Frontiers in Pharmacology 3.

52. Shemer, A., Scheyltjens, I., Frumer, G.R., Kim, J.-S., Grozovski, J., Ayanaw, S., Dassa, B., Hove, H.V., Chappell-Maor, L., Boura-Halfon, S., et al. (2020). Interleukin-10 Prevents Pathological Microglia Hyperactivation following Peripheral Endotoxin Challenge. Immunity 0. 10.1016/j.immuni.2020.09.018.

53. Luan, H.H., Wang, A., Hilliard, B.K., Carvalho, F., Rosen, C.E., Ahasic, A.M., Herzog, E.L., Kang, I., Pisani, M.A., Yu, S., et al. (2019). GDF15 Is an Inflammation-Induced Central Mediator of Tissue Tolerance. Cell 178, 1231–1244.e11. 10.1016/j.cell.2019.07.033.

54. Cho, H.-Y., Yang, Y.G., Jeon, Y., Lee, C.-K., Choi, I., and Lee, S.-W. (2021). VSIG4(+) peritoneal macrophages induce apoptosis of double-positive thymocyte via the secretion of TNF-α in a CLP-induced sepsis model resulting in thymic atrophy. Cell Death Dis 12, 1–14. 10.1038/s41419-021-03806-5.

55. Janosevic, D., Myslinski, J., McCarthy, T.W., Zollman, A., Syed, F., Xuei, X., Gao, H., Liu, Y.-L., Collins, K.S., Cheng, Y.-H., et al. (2021). The orchestrated cellular and molecular responses of the kidney to endotoxin define a precise sepsis timeline. eLife 10, e62270. 10.7554/eLife.62270.

56. Bartee, E., and McFadden, G. (2013). Cytokine synergy: An underappreciated contributor to innate anti-viral immunity. Cytokine 63, 237–240. 10.1016/j.cyto.2013.04.036.

57. Wong, G.H.W., and Goeddel, D.V. (1986). Tumour necrosis factors α and β inhibit virus replication and synergize with interferons. Nature 323, 819–822. 10.1038/323819a0.

58. Bevilacqua, M.P., Pober, J.S., Majeau, G.R., Fiers, W., Cotran, R.S., and Gimbrone, M.A. (1986). Recombinant tumor necrosis factor induces procoagulant activity in cultured human vascular endothelium: characterization and comparison with the actions of interleukin 1. PNAS 83, 4533–4537. 10.1073/pnas.83.12.4533.

59. Last-Barney, K., Homon, C.A., Faanes, R.B., and Merluzzi, V.J. (1988). Synergistic and overlapping activities of tumor necrosis factor-alpha and IL-1. The Journal of Immunology 141, 527–530.

60. Harigai, M., Hara, M., Kitani, A., Norioka, K., Hirose, T., Hirose, W., Suzuki, K., Kawakami, M., Masuda, K., and Shinmei, M. (1991). Interleukin 1 and tumor necrosis factor-alpha synergistically increase the production of interleukin 6 in human synovial fibroblast. J Clin Lab Immunol 34, 107–113.

61. Faquin, W.C., Schneider, T.J., and Goldberg, M.A. (1992). Effect of Inflammatory Cytokines on Hypoxia-Induced Erythropoietin Production. Blood 79, 1987–1994. 10.1182/blood.V79.8.1987.1987.

62. Selleri, C., Sato, T., Anderson, S., Young, N.S., and Maciejewski, J.P. (1995). Interferon-γ and tumor necrosis factor-α suppress both early and late stages of hematopoiesis and induce programmed cell death. Journal of Cellular Physiology 165, 538–546. 10.1002/jcp.1041650312.

63. Fish, S., Proujansky, R., and Reenstra, W. (1999). Synergistic effects of interferon γ and tumour necrosis factor α on T84 cell function. Gut 45, 191–198.

64. Dinarello, C.A., Cannon, J.G., Wolff, S.M., Bernheim, H.A., Beutler, B., Cerami, A., Figari, I.S., Palladino, M.A., Jr, and O’Connor, J.V. (1986). Tumor necrosis factor (cachectin) is an endogenous pyrogen and induces production of interleukin 1. Journal of Experimental Medicine 163, 1433–1450. 10.1084/jem.163.6.1433.

65. Gouwy, M., Struyf, S., Proost, P., and Van Damme, J. (2005). Synergy in cytokine and chemokine networks amplifies the inflammatory response. Cytokine & Growth Factor Reviews 16, 561–580. 10.1016/j.cytogfr.2005.03.005.

66. Doherty, G.M., Lange, J.R., Langstein, H.N., Alexander, H.R., Buresh, C.M., and Norton, J.A. (1992). Evidence for IFN-gamma as a mediator of the lethality of endotoxin and tumor necrosis factor-alpha. The Journal of Immunology 149, 1666–1670.

67. Suk, K., Kim, S., Kim, Y.-H., Kim, K.-A., Chang, I., Yagita, H., Shong, M., and Lee, M.-S. (2001). IFN-γ/TNF-α Synergism as the Final Effector in Autoimmune Diabetes: A Key Role for STAT1/IFN Regulatory Factor-1 Pathway in Pancreatic β Cell Death1. The Journal of Immunology 166, 4481–4489. 10.4049/jimmunol.166.7.4481.

68. Karki, R., Sharma, B.R., Tuladhar, S., Williams, E.P., Zalduondo, L., Samir, P., Zheng, M., Sundaram, B., Banoth, B., Malireddi, R.K.S., et al. (2021). Synergism of TNF-α and IFN-γ Triggers Inflammatory Cell Death, Tissue Damage, and Mortality in SARS-CoV-2 Infection and Cytokine Shock Syndromes. Cell 184, 149-168.e17. 10.1016/j.cell.2020.11.025.

69. Babaeijandaghi, F., Paiero, A., Long, R., Tung, L.W., Smith, S.P., Cheng, R., Smandych, J., Kajabadi, N., Chang, C.-K., Ghassemi, A., et al. (2022). TNFα and IFNγ cooperate for efficient pro- to anti-inflammatory transition of macrophages during muscle regeneration. Proceedings of the National Academy of Sciences 119, e2209976119. 10.1073/pnas.2209976119.

70. Okusawa, S., Gelfand, J.A., Ikejima, T., Connolly, R.J., and Dinarello, C.A. (1988). Interleukin 1 induces a shock-like state in rabbits. Synergism with tumor necrosis factor and the effect of cyclooxygenase inhibition. J Clin Invest 81, 1162–1172. 10.1172/JCI113431.

71. Waage, A., and Espevik, T. (1988). Interleukin 1 potentiates the lethal effect of tumor necrosis factor alpha/cachectin in mice. J Exp Med 167, 1987–1992. 10.1084/jem.167.6.1987.

72. Russell, D.A., Tucker, K.K., Chinookoswong, N., Thompson, R.C., and Kohno, T. (1995). Combined Inhibition of Interleukin-1 and Tumor Necrosis Factor in Rodent Endotoxemia: Improved Survival and Organ Function. The Journal of Infectious Diseases 171, 1528–1538. 10.1093/infdis/171.6.1528.

73. Jahnke, A., and P. Johnson, J. (1994). Synergistic activation of intercellular adhesion molecule 1 (ICAM-1) by TNF-α and IFN-γ is mediated by p65/p50 and p65/c-Rel and interferon-responsive factor Statlα (p91) that can be activated by both IFN-γ and IFN-α. FEBS Letters 354, 220–226. 10.1016/0014-5793(94)01130-3.

74. Pine, R. (1997). Convergence of TNFα and IFNγ signalling pathways through synergistic induction of IRF-1/ISGF-2 is mediated by a composite GAS/κB promoter element. Nucleic Acids Research 25, 4346–4354. 10.1093/nar/25.21.4346.

75. Carswell, E.A., Old, L.J., Kassel, R.L., Green, S., Fiore, N., and Williamson, B. (1975). An endotoxin-induced serum factor that causes necrosis of tumors. Proceedings of the National Academy of Sciences 72, 3666–3670. 10.1073/pnas.72.9.3666.

76. Clark, I.A. (2007). How TNF was recognized as a key mechanism of disease. Cytokine & Growth Factor Reviews 18, 335–343. 10.1016/j.cytogfr.2007.04.002.

77. Brenner, D., Blaser, H., and Mak, T.W. (2015). Regulation of tumour necrosis factor signalling: live or let die. Nat Rev Immunol 15, 362–374. 10.1038/nri3834.

78. Waage, A., Halstensen, A., and Espevik, T. (1987). ASSOCIATION BETWEEN TUMOUR NECROSIS FACTOR IN SERUM AND FATAL OUTCOME IN PATIENTS WITH MENINGOCOCCAL DISEASE. The Lancet 329, 355–357. 10.1016/S0140-6736(87)91728-4.

79. DeForge, L.E., Nguyen, D.T., Kunkel, S.L., and Remick, D.G. (1990). Regulation of the pathophysiology of tumor necrosis factor. J Lab Clin Med 116, 429–438.

80. Tracey, K.J., Fong, Y., Hesse, D.G., Manogue, K.R., Lee, A.T., Kuo, G.C., Lowry, S.F., and Cerami, A. (1987). Anti-cachectin/TNF monoclonal antibodies prevent septic shock during lethal bacteraemia. Nature 330, 662–664. 10.1038/330662a0.

81. Beutler, B., Milsark, I.W., and Cerami, A.C. (1985). Passive Immunization Against Cachectin/Tumor Necrosis Factor Protects Mice from Lethal Effect of Endotoxin. Science 229, 869–871. 10.1126/science.3895437.

82. Fisher, C.J., Agosti, J.M., Opal, S.M., Lowry, S.F., Balk, R.A., Sadoff, J.C., Abraham, E., Schein, R.M.H., and Benjamin, E. (1996). Treatment of Septic Shock with the Tumor Necrosis Factor Receptor:Fc Fusion Protein. N Engl J Med 334, 1697–1702. 10.1056/NEJM199606273342603.

83. Angus, D.C. (2011). Management of Sepsis: A 47-Year-Old Woman With an Indwelling Intravenous Catheter and Sepsis. JAMA 305, 1469–1477. 10.1001/jama.2011.438.

84. van der Poll, T., and Opal, S.M. (2008). Host–pathogen interactions in sepsis. The Lancet Infectious Diseases 8, 32–43. 10.1016/S1473-3099(07)70265-7.

85. Brown, K.A., Brown, G.A., Lewis, S.M., Beale, R., and Treacher, D.F. (2016). Targeting cytokines as a treatment for patients with sepsis: A lost cause or a strategy still worthy of pursuit? International Immunopharmacology 36, 291–299. 10.1016/j.intimp.2016.04.041.

86. Fekade, D., Knox, K., Hussein, K., Melka, A., Lalloo, D.G., Coxon, R.E., and Warrell, D.A. (1996). Prevention of Jarisch–Herxheimer Reactions by Treatment with Antibodies against Tumor Necrosis Factor α. N Engl J Med 335, 311–315. 10.1056/NEJM199608013350503.

87. Coxon, R.E., Fekade, D., Knox, K., Hussein, K., Melka, A., Daniel, A., Griffin, G.G., and Warrell, D.A. (1997). The effect of antibody against TNF alpha on cytokine response in Jarisch-Herxheimer reactions of louse-borne relapsing fever. QJM: An International Journal of Medicine 90, 213–221. 10.1093/qjmed/90.3.213.

88. Steinmetz, T., Schaadt, M., Gahl, R., Schenk, V., Diehl, V., and Pfreundschuh, M. (1988). Phase I Study of 24-Hour Continuous Intravenous Infusion of Recombinant Human Tumor Necrosis Factor. Journal of Biological Response Modifiers 7, 417–423.

89. Schett, G., Elewaut, D., McInnes, I.B., Dayer, J.-M., and Neurath, M.F. (2013). How Cytokine Networks Fuel Inflammation: Toward a cytokine-based disease taxonomy. Nat Med 19, 822–824. 10.1038/nm.3260.

90. Schett, G., McInnes, I.B., and Neurath, M.F. (2021). Reframing Immune-Mediated Inflammatory Diseases through Signature Cytokine Hubs. N Engl J Med 385, 628–639. 10.1056/NEJMra1909094.

91. Dore, E., and Boilard, E. (2019). Roles of secreted phospholipase A2 group IIA in inflammation and host defense. Biochimica et Biophysica Acta (BBA) - Molecular and Cell Biology of Lipids 1864, 789–802. 10.1016/j.bbalip.2018.08.017.

92. Dennis, E.A., and Norris, P.C. (2015). Eicosanoid storm in infection and inflammation. Nat Rev Immunol 15, 511–523. 10.1038/nri3859.

93. Duchez, A.-C., Boudreau, L.H., Naika, G.S., Bollinger, J., Belleannée, C., Cloutier, N., Laffont, B., Mendoza-Villarroel, R.E., Lévesque, T., Rollet-Labelle, E., et al. (2015). Platelet microparticles are internalized in neutrophils via the concerted activity of 12-lipoxygenase and secreted phospholipase A2-IIA. Proceedings of the National Academy of Sciences 112, E3564–E3573. 10.1073/pnas.1507905112.

94. Neidlinger, N.A., Larkin, S.K., Bhagat, A., Victorino, G.P., and Kuypers, F.A. (2006). Hydrolysis of Phosphatidylserine-exposing Red Blood Cells by Secretory Phospholipase A2 Generates Lysophosphatidic Acid and Results in Vascular Dysfunction*. Journal of Biological Chemistry 281, 775–781. 10.1074/jbc.M505790200.

95. Murakami, M., Taketomi, Y., Miki, Y., Sato, H., Hirabayashi, T., and Yamamoto, K. (2011). Recent progress in phospholipase A2 research: From cells to animals to humans. Progress in Lipid Research 50, 152–192. 10.1016/j.plipres.2010.12.001.

96. Watanabe, K., Taketomi, Y., Miki, Y., Kugiyama, K., and Murakami, M. (2020). Group V secreted phospholipase A2 plays a protective role against aortic dissection. J. Biol. Chem. 295, 10092–10111. 10.1074/jbc.RA120.013753.

97. Miki, Y., Taketomi, Y., Kidoguchi, Y., Yamamoto, K., Muramatsu, K., Nishito, Y., Park, J., Hosomi, K., Mizuguchi, K., Kunisawa, J., et al. (2022). Group IIA secreted phospholipase A_2_ controls skin carcinogenesis and psoriasis by shaping the gut microbiota. JCI Insight 7. 10.1172/jci.insight.152611.

98. Seifert, S.A., Armitage, J.O., and Sanchez, E.E. (2022). Snake Envenomation. N Engl J Med 386, 68–78. 10.1056/NEJMra2105228.

99. Arce-Bejarano, R., Lomonte, B., and Gutiérrez, J.M. (2014). Intravascular hemolysis induced by the venom of the Eastern coral snake, Micrurus fulvius, in a mouse model: Identification of directly hemolytic phospholipases A2. Toxicon 90, 26–35. 10.1016/j.toxicon.2014.07.010.

100. Fay, K.T., Ford, M.L., and Coopersmith, C.M. (2017). The intestinal microenvironment in sepsis. Biochimica et Biophysica Acta (BBA) - Molecular Basis of Disease 1863, 2574– 2583. 10.1016/j.bbadis.2017.03.005.

101. Haussner, F., Chakraborty, S., Halbgebauer, R., and Huber-Lang, M. (2019). Challenge to the Intestinal Mucosa During Sepsis. Frontiers in Immunology 10.

102. Htwe, Y.M., Wang, H., Belvitch, P., Meliton, L., Bandela, M., Letsiou, E., and Dudek, S.M. (2021). Group V Phospholipase A2 Mediates Endothelial Dysfunction and Acute Lung Injury Caused by Methicillin-Resistant Staphylococcus Aureus. Cells 10, 1731. 10.3390/cells10071731.

103. Muñoz, N.M., Meliton, A.Y., Meliton, L.N., Dudek, S.M., and Leff, A.R. (2009). Secretory group V phospholipase A2 regulates acute lung injury and neutrophilic inflammation caused by LPS in mice. American Journal of Physiology-Lung Cellular and Molecular Physiology 296, L879–L887. 10.1152/ajplung.90580.2008.

104. Soumillon, M., Cacchiarelli, D., Semrau, S., van Oudenaarden, A., and Mikkelsen, T.S. (2014). Characterization of directed differentiation by high-throughput single-cell RNA-Seq. bioRxiv. 10.1101/003236.

105. Yamamoto, K., Miki, Y., Sato, H., Murase, R., Taketomi, Y., and Murakami, M. (2017). Chapter Five - Secreted Phospholipase A2 Specificity on Natural Membrane Phospholipids. In Methods in Enzymology Enzymology at the Membrane Interface: Interfacial Enzymology and Protein-Membrane Binding., M. H. Gelb, ed. (Academic Press), pp. 101–117. 10.1016/bs.mie.2016.09.007.

106. Bligh, E.G., and Dyer, W.J. (1959). A rapid method of total lipid extraction and purification. Can J Biochem Physiol 37, 911–917. 10.1139/o59-099.

107. Yamamoto, K., Miki, Y., Sato, M., Taketomi, Y., Nishito, Y., Taya, C., Muramatsu, K., Ikeda, K., Nakanishi, H., Taguchi, R., et al. (2015). The role of group IIF-secreted phospholipase A2 in epidermal homeostasis and hyperplasia. Journal of Experimental Medicine 212, 1901–1919. 10.1084/jem.20141904.

108. Islam, S., Zeisel, A., Joost, S., La Manno, G., Zajac, P., Kasper, M., Lönnerberg, P., and Linnarsson, S. (2014). Quantitative single-cell RNA-seq with unique molecular identifiers. Nat Methods 11, 163–166. 10.1038/nmeth.2772.

109. Srivastava, A., Sarkar, H., Gupta, N., and Patro, R. (2016). RapMap: a rapid, sensitive and accurate tool for mapping RNA-seq reads to transcriptomes. Bioinformatics 32, i192– i200. 10.1093/bioinformatics/btw277.

110. García-Alcalde, F., Okonechnikov, K., Carbonell, J., Cruz, L.M., Götz, S., Tarazona, S., Dopazo, J., Meyer, T.F., and Conesa, A. (2012). Qualimap: evaluating next-generation sequencing alignment data. Bioinformatics 28, 2678–2679. 10.1093/bioinformatics/bts503.

111. Ewels, P., Magnusson, M., Lundin, S., and Käller, M. (2016). MultiQC: summarize analysis results for multiple tools and samples in a single report. Bioinformatics 32, 3047– 3048. 10.1093/bioinformatics/btw354.

112. Robinson, M.D., McCarthy, D.J., and Smyth, G.K. (2010). edgeR: a Bioconductor package for differential expression analysis of digital gene expression data. Bioinformatics 26, 139–140. 10.1093/bioinformatics/btp616.

113. Huang, D.W., Sherman, B.T., and Lempicki, R.A. (2009). Bioinformatics enrichment tools: paths toward the comprehensive functional analysis of large gene lists. Nucleic Acids Research 37, 1–13. 10.1093/nar/gkn923.

114. Huang, D.W., Sherman, B.T., and Lempicki, R.A. (2009). Systematic and integrative analysis of large gene lists using DAVID bioinformatics resources. Nat Protoc 4, 44–57. 10.1038/nprot.2008.211.

115. Gu, Z., Eils, R., and Schlesner, M. (2016). Complex heatmaps reveal patterns and correlations in multidimensional genomic data. Bioinformatics 32, 2847–2849. 10.1093/bioinformatics/btw313.

116. Gu, Z., Gu, L., Eils, R., Schlesner, M., and Brors, B. (2014). circlize implements and enhances circular visualization in R. Bioinformatics 30, 2811–2812. 10.1093/bioinformatics/btu393.

117. Carbonetto, P., Sarkar, A., Wang, Z., and Stephens, M. (2022). Non-negative matrix factorization algorithms greatly improve topic model fits. 10.48550/arXiv.2105.13440.

118. Hien, L.T.K., and Gillis, N. (2021). Algorithms for Nonnegative Matrix Factorization with the Kullback–Leibler Divergence. J Sci Comput 87, 93. 10.1007/s10915-021-01504-0.

119. van der Maaten, L., and Hinton, G. (2008). Visualizing Data using t-SNE. Journal of Machine Learning Research 9, 2579–2605.

120. Rosenberg, N.A., Pritchard, J.K., Weber, J.L., Cann, H.M., Kidd, K.K., Zhivotovsky, L.A., and Feldman, M.W. (2002). Genetic Structure of Human Populations. Science 298, 2381–2385. 10.1126/science.1078311.

121. Soneson, C., and Robinson, M.D. (2018). Bias, robustness and scalability in single-cell differential expression analysis. Nat Methods 15, 255–261. 10.1038/nmeth.4612.

122. Wang, T., Li, B., Nelson, C.E., and Nabavi, S. (2019). Comparative analysis of differential gene expression analysis tools for single-cell RNA sequencing data. BMC Bioinformatics 20, 40. 10.1186/s12859-019-2599-6.

123. Stephens, M. (2017). False discovery rates: a new deal. Biostatistics 18, 275–294. 10.1093/biostatistics/kxw041.

124. Geer, L.Y., Marchler-Bauer, A., Geer, R.C., Han, L., He, J., He, S., Liu, C., Shi, W., and Bryant, S.H. (2010). The NCBI BioSystems database. Nucleic Acids Research 38, D492– D496. 10.1093/nar/gkp858.

125. Cerami, E.G., Gross, B.E., Demir, E., Rodchenkov, I., Babur, Ö., Anwar, N., Schultz, N., Bader, G.D., and Sander, C. (2011). Pathway Commons, a web resource for biological pathway data. Nucleic Acids Research 39, D685–D690. 10.1093/nar/gkq1039.

126. Rodchenkov, I., Babur, O., Luna, A., Aksoy, B.A., Wong, J.V., Fong, D., Franz, M., Siper, M.C., Cheung, M., Wrana, M., et al. (2020). Pathway Commons 2019 Update: integration, analysis and exploration of pathway data. Nucleic Acids Research 48, D489– D497. 10.1093/nar/gkz946.

127. Subramanian, A., Tamayo, P., Mootha, V.K., Mukherjee, S., Ebert, B.L., Gillette, M.A., Paulovich, A., Pomeroy, S.L., Golub, T.R., Lander, E.S., et al. (2005). Gene set enrichment analysis: A knowledge-based approach for interpreting genome-wide expression profiles. Proceedings of the National Academy of Sciences 102, 15545–15550. 10.1073/pnas.0506580102.

128. Liberzon, A., Subramanian, A., Pinchback, R., Thorvaldsdóttir, H., Tamayo, P., and Mesirov, J.P. (2011). Molecular signatures database (MSigDB) 3.0. Bioinformatics 27, 1739– 1740. 10.1093/bioinformatics/btr260.

129. Liberzon, A., Birger, C., Thorvaldsdóttir, H., Ghandi, M., Mesirov, J.P., and Tamayo, P. (2015). The Molecular Signatures Database Hallmark Gene Set Collection. Cell Systems 1, 417–425. 10.1016/j.cels.2015.12.004.

130. Ashburner, M., Ball, C.A., Blake, J.A., Botstein, D., Butler, H., Cherry, J.M., Davis, A.P., Dolinski, K., Dwight, S.S., Eppig, J.T., et al. (2000). Gene Ontology: tool for the unification of biology. Nat Genet 25, 25–29. 10.1038/75556.

131. The Gene Ontology Consortium (2021). The Gene Ontology resource: enriching a GOld mine. Nucleic Acids Research 49, D325–D334. 10.1093/nar/gkaa1113.

132. Wang, G., Sarkar, A., Carbonetto, P., and Stephens, M. (2020). A simple new approach to variable selection in regression, with application to genetic fine mapping. Journal of the Royal Statistical Society: Series B (Statistical Methodology) 82, 1273–1300. 10.1111/rssb.12388.

133. Schrode, N., Seah, C., Deans, P.J.M., Hoffman, G., and Brennand, K.J. (2021). Analysis framework and experimental design for evaluating synergy-driving gene expression. Nat Protoc 16, 812–840. 10.1038/s41596-020-00436-7.

134. Ritchie, M.E., Phipson, B., Wu, D., Hu, Y., Law, C.W., Shi, W., and Smyth, G.K. (2015). limma powers differential expression analyses for RNA-sequencing and microarray studies. Nucleic Acids Research 43, e47. 10.1093/nar/gkv007.

135. Patil, A., and Patil, A. (2022). CellKb Immune: a manually curated database of mammalian hematopoietic marker gene sets for rapid cell type identification. bioRxiv. 10.1101/2020.12.01.389890.

136. Single-cell transcriptomics of 20 mouse organs creates a Tabula Muris (2018). Nature 562, 367–372. 10.1038/s41586-018-0590-4.

137. Uhlén, M., Fagerberg, L., Hallström, B.M., Lindskog, C., Oksvold, P., Mardinoglu, A., Sivertsson, Å., Kampf, C., Sjöstedt, E., Asplund, A., et al. (2015). Tissue-based map of the human proteome. Science 347, 1260419. 10.1126/science.1260419.

138. Diehl, A.D., Meehan, T.F., Bradford, Y.M., Brush, M.H., Dahdul, W.M., Dougall, D.S., He, Y., Osumi-Sutherland, D., Ruttenberg, A., Sarntivijai, S., et al. (2016). The Cell Ontology 2016: enhanced content, modularization, and ontology interoperability. Journal of Biomedical Semantics 7, 44. 10.1186/s13326-016-0088-7.

139. Mungall, C.J., Torniai, C., Gkoutos, G.V., Lewis, S.E., and Haendel, M.A. (2012). Uberon, an integrative multi-species anatomy ontology. Genome Biology 13, R5. 10.1186/gb-2012-13-1-r5.

140. Schriml, L.M., Arze, C., Nadendla, S., Chang, Y.-W.W., Mazaitis, M., Felix, V., Feng, G., and Kibbe, W.A. (2012). Disease Ontology: a backbone for disease semantic integration. Nucleic Acids Research 40, D940–D946. 10.1093/nar/gkr972.

141. Wolf, F.A., Angerer, P., and Theis, F.J. (2018). SCANPY: large-scale single-cell gene expression data analysis. Genome Biology 19, 15. 10.1186/s13059-017-1382-0.

